# Using all gene families vastly expands data available for phylogenomic inference

**DOI:** 10.1101/2021.09.22.461252

**Authors:** Megan L. Smith, Dan Vanderpool, Matthew W. Hahn

## Abstract

Traditionally, single-copy orthologs have been the gold standard in phylogenomics. Most phylogenomic studies identify putative single-copy orthologs using clustering approaches and retain families with a single sequence per species. This limits the amount of data available by excluding larger families. Recent advances have suggested several ways to include data from larger families. For instance, tree-based decomposition methods facilitate the extraction of orthologs from large families. Additionally, several methods for species tree inference are robust to the inclusion of paralogs, and could use all of the data from larger families. Here, we explore the effects of using all families for phylogenetic inference by examining relationships among 26 primate species in detail, and by analyzing five additional datasets. We compare single-copy families, orthologs extracted using tree-based decomposition approaches, and all families with all data. We explore several species tree inference methods, finding that identical trees are returned across nearly all subsets of the data and methods for primates. The relationships among Platyrrhini remain contentious; however, the species tree inference method matters more than the subset of data used. Using data from larger gene families drastically increases the number of genes available and leads to consistent estimates of branch lengths, nodal certainty and concordance, and inferences of introgression in primates. For the other datasets, topological inferences are consistent whether single-copy families or orthologs extracted using decomposition approaches are analyzed. Using larger gene families is a promising approach to include more data in phylogenomics without sacrificing accuracy, at least when high-quality genomes are available.

## Introduction

Advances in sequencing technology have led to the availability of more genomic data than ever before, and the promise of phylogenomics is the application of this data to infer species relationships (Scornavacca et al. 2020). Essential to the application of genomic data to phylogenetic inference is the identification of homologous genes, or genes that share a common ancestor. Homologous genes may share a common ancestor due to speciation (orthologs) or duplication (paralogs). Since the terms ortholog and paralog were coined (Fitch 1970), orthologs have been considered the appropriate genes for phylogenetic inference because they are related only through speciation events, and therefore are thought to best reflect species relationships. Thus, identifying orthologs is a central part of most phylogenomic pipelines.

Nearly all pipelines for extracting putative orthologs from genomic data begin with a clustering step (Figure 1). Clustering approaches aim to identify sets of homologous genes. While the details vary, these approaches generally begin with pairwise comparisons of all sequences across genomes, identify putative pairwise homologs, and then use clustering approaches to attempt to group many sets of these genes together (reviewed in Altenhoff et al. 2019). The end-products of graph-based clustering approaches are clusters of orthologs and paralogs—i.e., gene families. Since most phylogenetic methods were designed for use with orthologs (and a single sequence per taxon), these groups must be further processed for downstream phylogenetic inference.

**Figure 1.**
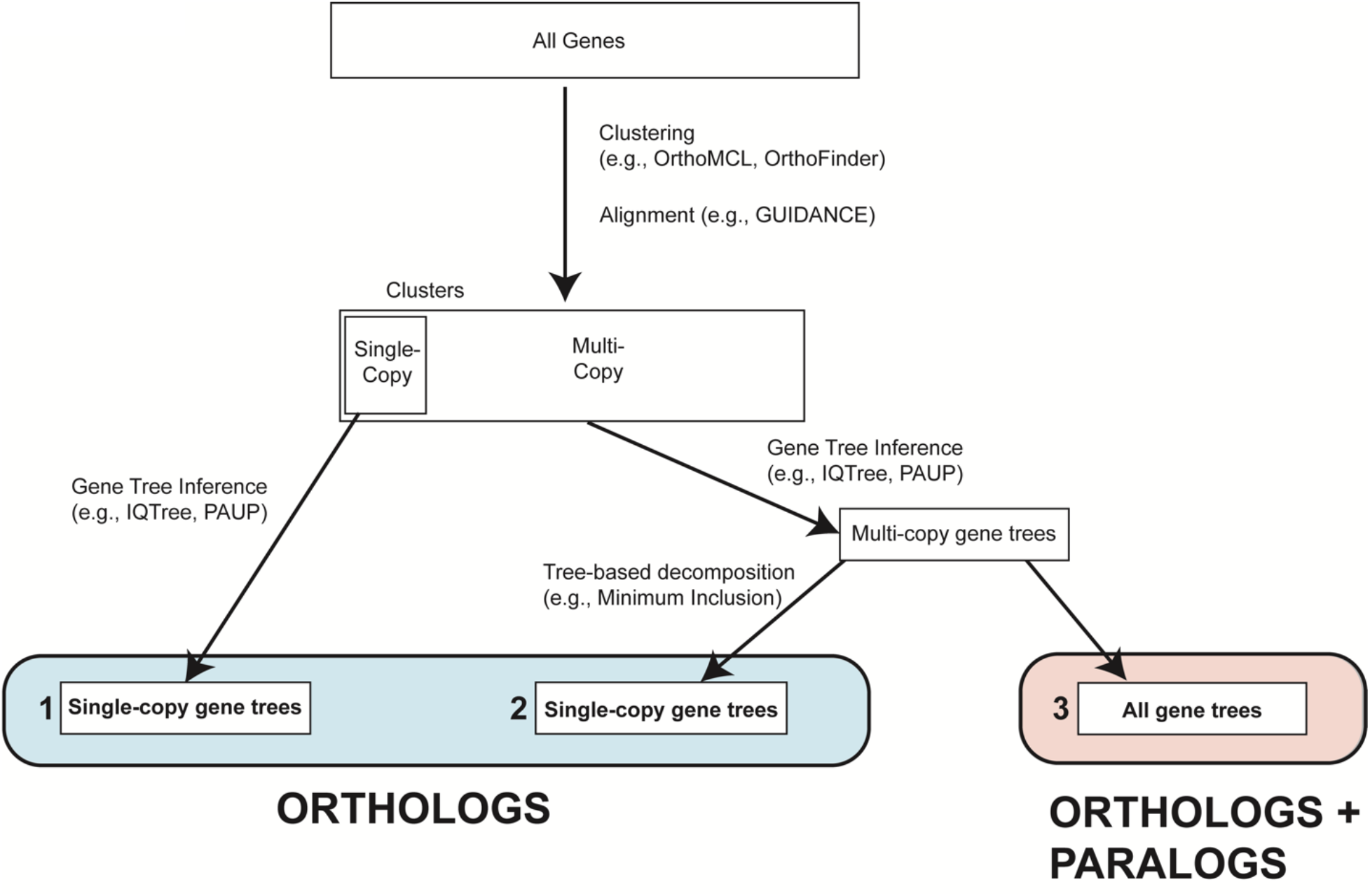
Conceptual overview of methods for inferring species trees from genomic data. We begin with All Genes, clustering them into gene families. We can then use single-copy ortholog clusters for inference (Dataset 1), use tree-based decomposition approaches to extract orthologs from all clusters (Dataset 2), or infer species trees from all clusters (i.e., from datasets including orthologs and paralogs; Dataset 3).

Three primary approaches have been used to process families for downstream inference (Figure 1; Step 1). The first and most common is to extract clusters with only a single copy in each species—these represent putative single-copy orthologs. Using single-copy families is generally seen as a conservative approach in phylogenomics, as these genes are likely to be orthologs; this choice also limits the amount of further downstream processing needed. However, the number of genes that are single-copy in all sampled species decreases sharply as additional species are included in the analyses (Emms and Kelly 2018), limiting the usefulness of this approach in many phylogenetic contexts.

In lieu of relying only on single-copy clusters, tree-based decomposition approaches for orthology detection can be applied to extract orthologous genes from clusters that may have more than one copy in one or more species (Figure 1; Step 2). Tree-based decomposition approaches attempt to infer whether nodes in gene trees represent duplication or speciation events, followed by the extraction of orthologs based on these node labels (reviewed in Altenhoff et al. 2019). Early tree-based approaches relied on gene tree reconciliation to a known species tree (e.g., Goodman et al. 1979), limiting their utility in cases where the species tree is unknown or uncertain. However, recent approaches have relaxed these requirements. For example, the method LOFT relies on a species overlap approach to identify duplication nodes in gene trees (van der Heijden et al. 2007). Similarly, the software package Agalma (Dunn et al. 2013), the methods of Yang and Smith (2014), and the new method, DISCO (Willson et al. 2021), all extract subtrees without duplicates to generate sets of orthologs. While the exact implementations vary, in general tree-based decomposition approaches aim to extract orthologous genes from families of any size. Tree-based approaches allow researchers to vastly increase the number of genes retained compared to using only the single-copy clusters. However, these approaches require that users construct gene trees and perform ortholog extraction for each gene family. Since gene trees must be constructed for all gene families, and some of these gene families may be rather large, these approaches can be substantially more computationally intensive than relying on single-copy clusters alone (Figure 1).

Finally, families containing both orthologs and paralogs could be used for phylogenetic inference. Although orthologs have traditionally been considered the appropriate genes for phylogenetics, methods for estimating phylogenies from data including paralogs were introduced more than forty years ago (Goodman et al. 1979; reviewed in Smith and Hahn 2021a). Recently, several popular methods for species tree estimation have been shown to be robust to the presence of paralogs (Hill et al. 2020; Legried et al. 2020; Markin and Eulenstein 2020; Yan et al. 2021). Of particular interest, quartet-based methods, such as ASTRAL (Zhang et al. 2018), should be robust to the inclusion of paralogs because the most common quartet is still expected to match the species tree even in the presence of gene duplication and loss. Given that all ortholog extraction methods may erroneously lead to the inclusion of paralogs, using methods that are robust to their inclusion is likely a good strategy no matter the method employed to process the output of clustering methods.

Though there have been several empirical comparisons between ortholog-detection methods (e.g., Fernández et al. 2018; Kallal et al. 2018; Altenhoff et al. 2019), along with several simulation-based (e.g., Legried et al. 2020; Zhang et al. 2020; Yan et al. 2021; Morel et al. 2022) and empirical (e.g., Yan et al. 2021) studies evaluating the effects of paralog inclusion on phylogenetic inference, several questions remain. First, a comparison of inference on single-copy clusters to tree-based decomposition methods and methods that use all of the data (i.e., use orthologs and paralogs for phylogenetic inference) would shed light on the advantages of the three approaches. In addition, the joint effects of dataset, missing data requirements, and gene and species tree inference method on species tree topology will provide information on the importance of each. Finally, questions remain about the effects of the dataset used on branch length estimates, measures of nodal support, and tests for introgression.

To address these questions, we focus our analysis on a recently published phylogenomic dataset that includes 26 species of primates and 3 outgroups (Vanderpool et al. 2020). The data consist of whole genomes from all 29 species. In the original study, Vanderpool et al. restricted inference to 1,730 single-copy clusters present in 27 of the 29 studied species, a relatively small proportion of the >20,000 genes available from each species; the species tree was inferred using concatenated maximum likelihood, concatenated maximum parsimony, and quartet-based approaches applied to gene trees inferred using both maximum likelihood and maximum parsimony. The authors found robust relationships among all species except the Platyrrhini (“New World Monkeys”), for which inferences differed across species-tree and gene-tree inference methods. In this paper, we compare inferences from three major subsets of the data: single-copy families, orthologs extracted from larger families using tree-based decomposition approaches, and all families including all data (orthologs + paralogs). These datasets are then compared in three different phylogenetic applications. First, we compare the species trees inferred from these datasets using several methods, including concatenation-based and gene-tree-based approaches. Second, we compare several measures of nodal support and nodal consistency, as well as branch length estimates across datasets. Finally, we perform tests of introgression and compare results across different datasets. In addition to analyzing the primate dataset, we assembled datasets from five different groups (two fungi datasets, one plant dataset, and two vertebrate datasets; Morel et al. 2022; Rasmussen and Kellis 2012), and compared species trees inferred from single-copy families, orthologs extracted from larger families using decomposition approaches, and all families for each. Our results suggest minimal effects of the subset of data used on downstream phylogenetic inference, while highlighting the fact that both tree-based decomposition approaches and approaches using both orthologs and paralogs greatly expand the amount of data available.

## Results

### Using all gene families vastly expands the data available for phylogenetics in primates

We compared three types of datasets produced by clustering approaches: single-copy clusters, orthologs extracted from all clusters using tree-based decomposition approaches, and all clusters (orthologs + paralogs) (Figure 1). For all datasets we considered both a stringent missing-data threshold (only those genes present in at least 27 of the 29 sampled species; MIN27) and a relaxed missing-data threshold (only those genes present in at least 4 of the 29 sampled species; MIN4). Gene duplication and loss appears to have had a substantial impact on these data. For example, the 11,555 gene families sampled in 27 of 29 species included 428,129 gene copies (an average of 37 gene copies per gene family), and only a small fraction of these genes (1820) were present in only a single copy in all sampled species. This suggests that most gene families studied here have experienced gene duplication and loss events during the evolutionary history of the primates. The first subset of the data considers only those clusters that included a single gene from each species (single-copy clusters; SCCs). While these genes are not guaranteed to be orthologs—due to the potential inclusion of pseudoorthologs (Doolittle and Brown 1994; Koonin 2005)—this is considered a safe approach and is often employed in phylogenomics. As expected, this dataset included the fewest genes (Table 1).

**Table 1:** Number of primate genes trees and gene copies included with different filtering approaches. LSD and TSD indicate when lineage-specific and both lineage-specific and two-species specific duplicates were trimmed; the subtree extraction method trims these automatically. The MIN4 dataset required a minimum of 4 taxa (out of 29 total), while the MIN27 dataset required a minimum of 27 taxa.

| Filter | MIN4 |  | MIN27 |  |
| --- | --- | --- | --- | --- |
|  | Gene families | Gene copies | Gene families | Gene copies |
| *Single-copy clusters* | 5771 | 94994 | 1820 | 51733 |
| *Lineage-specific duplicates (LSD)* | 13627 | 297831 | 7693 | 219441 |
| *Two-species duplicates (TSD)* | 14931 | 332718 | 8719 | 248759 |
| *Maximum Inclusion* | 27880 | 331990 | 4849 | 137733 |
| *Maximum Inclusion (LSD)* | 22360 | 464224 | 11479 | 327434 |
| *Maximum Inclusion (TSD)* | 21793 | 473000 | 12046 | 343652 |
| *Monophyletic Outgroups* | 9724 | 200503 | 4805 | 136749 |
| *Monophyletic Outgroups (LSD)* | 16962 | 387915 | 10222 | 291374 |
| *Monophyletic Outgroups (TSD)* | 17104 | 390584 | 10254 | 292257 |
| *Subtree Extraction* | 20562 | 470465 | 12198 | 347994 |
| *All Paralogs* | 18484 | 568342 | 11555 | 454509 |
| *One Paralogs* | 18484 | 428129 | 11555 | 330115 |

Tree-based decomposition approaches aim to extract orthologous genes from any cluster/family. We constructed gene trees for all clusters, and then used several tree-based approaches to extract orthologous genes. First, we considered those clusters in which all duplications were specific to a single lineage and kept a single gene-copy from this lineage. When duplications are restricted to a single lineage, choosing one of the copies as the ortholog cannot mislead phylogenetic inference regardless of which sequence is retained (see Figure 1d from Smith and Hahn 2021a; Figure S1a). This dataset (“lineage-specific duplicates”; LSDs) included more than 4X as many genes as the SCC dataset (Table 1). Next, we further expanded our criteria to include those clusters with duplications specific to a pair of lineages (“two-species duplicates”; TSDs; Figure S1b). Such duplications also cannot mislead topological inference, though picking a non-orthologous pair could lead to longer branches. It is straightforward to pick the most closely related pair of genes from the two species, which should not mislead either topological or branch length inferences; including these genes further expanded the dataset compared to the LSD dataset (Table 1).

We considered two tree-based decomposition approaches from Yang and Smith (2014): maximum inclusion (MI) and monophyletic outgroups (MO). The MI approach takes a gene tree and iteratively extracts subtrees with the highest number of taxa without taxon duplication, until it cannot extract anymore subtrees with the minimum number of taxa. The MO approach considers only those gene trees with a monophyletic outgroup, roots the tree, and infers gene duplications from the root to the tips, pruning at nodes with duplications. These two approaches were each applied to three datasets: the original gene trees, the original gene trees trimmed to remove lineage-specific duplicates, and the original gene trees trimmed to remove both lineage-specific and two-species duplicates. We explored the effects of additional filtering and alternative parameters for the MI approach; as these changes had minimal effects, the results are presented in the Supplementary Material (Appendix A). We also considered a new tree-based decomposition approach: subtree extraction (SE). In this approach, we midpoint-root gene trees, trimming away lineage-specific and two-species duplicates. We then extract subtrees that include a single representative from each taxon (i.e., subtrees with no duplicates) and keep those trees that meet minimum taxon-sampling thresholds (Figure S1c, d).

All tree-based approaches further expanded the amount of data available (Table 1). Since the SE and MI approaches are highly similar (neither requires an outgroup, and both aim to extract subtrees with no duplication events), we further examined the genes extracted using the two approaches. We compared the MI dataset with TSDs trimmed and a minimum of 27 taxa to the SE dataset with a minimum of 27 taxa sampled (this method trims TSDs internally). The number of trees extracted using the two approaches was very similar (12,046 vs 12,198 genes in the MI and SE datasets, respectively). For the 12,046 trees in the MI dataset, there was no analog in the SE dataset for 2.4%, there was an identical tree in the SE dataset for 92.7%, and there was a similar tree in the SE dataset for 4.8% (median Robinson-Foulds distance of these trees=2.0). Thus, the MI and SE approaches extract very similar subsets of trees from the original clusters.

Finally, we considered two approaches that made no attempt to remove paralogs from the dataset. We considered one dataset in which all orthologs and paralogs were included (“All Paralogs”). This dataset was the most complete, as, even though it had fewer gene trees than some tree-based approaches, the gene trees from these tree-based approaches are subtrees extracted from this full dataset. Therefore, this dataset includes the most gene copies (Table 1). This dataset cannot be analyzed using concatenation methods because these approaches require an alignment that includes a single sequence for each species. To address this, and to evaluate the effects of stochastic sampling of paralogs, we also included a dataset in which a single gene (without regard to whether it was an ortholog or paralog) was sampled at random from each species (“One Paralogs”).

In total, we considered 20 subsets of the data each with MIN4 and MIN27 taxon sampling. The number of gene families ranged from 1,820 to 27,900, and the number of gene copies ranged from 51,773 to 568,342 (Table 1). Clearly, considering only SCCs drastically restricts the amount of data available, in terms of the number of gene trees (Table 1), the number of gene copies (Table 1), the number of decisive sites for each branch of the species tree (Figure 2A), and the number of gene trees informative about each branch of the species tree (Figure 2B). All other datasets are subsets of the All Paralogs dataset, and thus this dataset is necessarily the most informative. Apart from the All Paralogs dataset, including a randomly sampled paralog (One Paralogs) leads to the most decisive sites (Figure 2A), though they are not necessarily the most accurate sites (see below and Figure 3). MI and SE lead to the most informative gene trees (Figure 2B).

**Figure 2.**
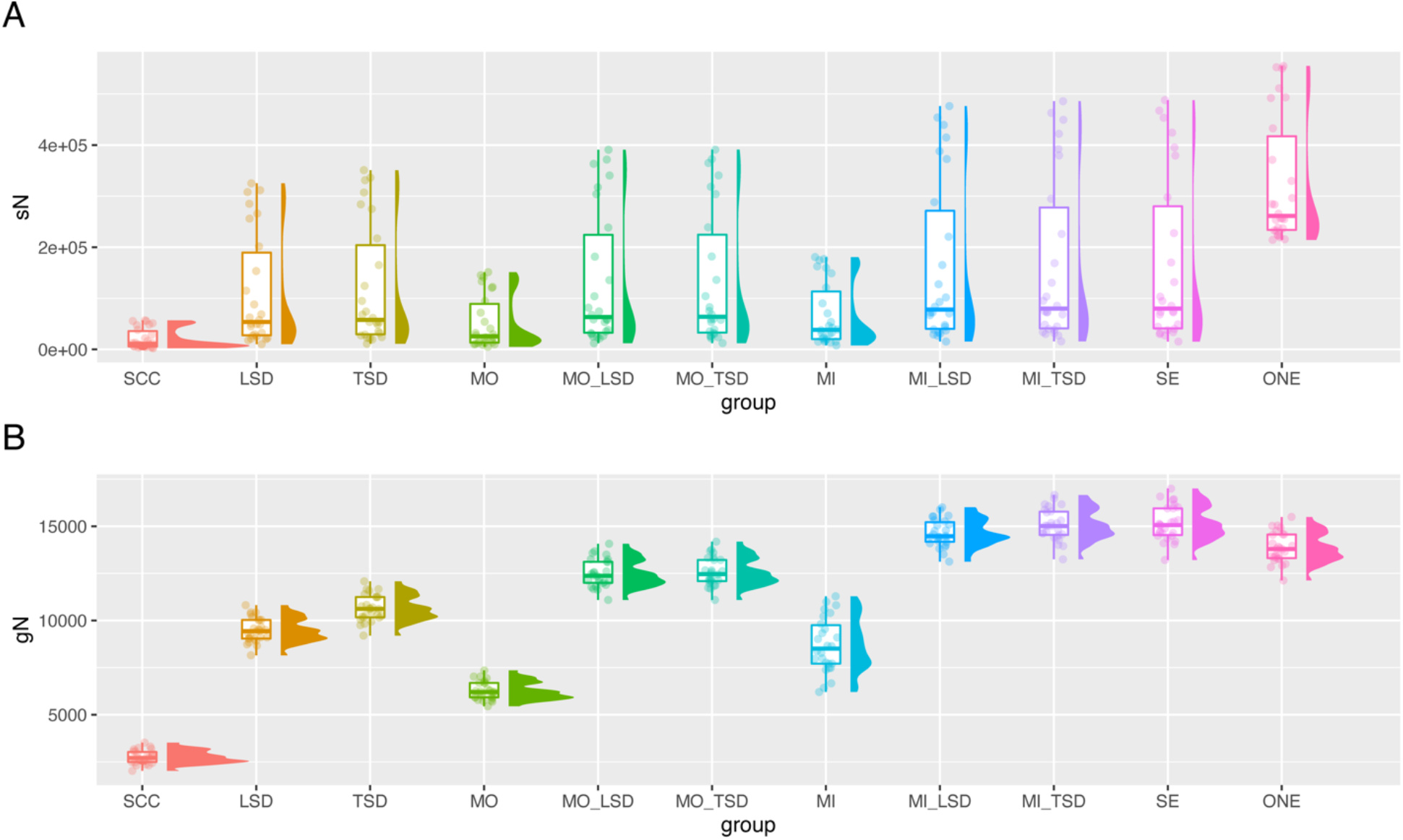
Numbers of informative genes and sites across datasets using the primate MIN27 datasets. A) The distribution of the number of decisive sites (across branches) as calculated in IQ-Tree. Decisive sites are defined in Minh et al. (2020). B) The distribution of the number of decisive gene trees (across branches) as calculated in IQ-Tree. Decisive gene trees are defined in Minh et al. (2020). SCC=single-copy clusters; LSD=lineage-specific duplicates; TSD=two-species duplicates; MO=monophyletic outgroup; MI=maximum inclusion; SE=subtree extraction; ONE=one paralogs.

**Figure 3.**
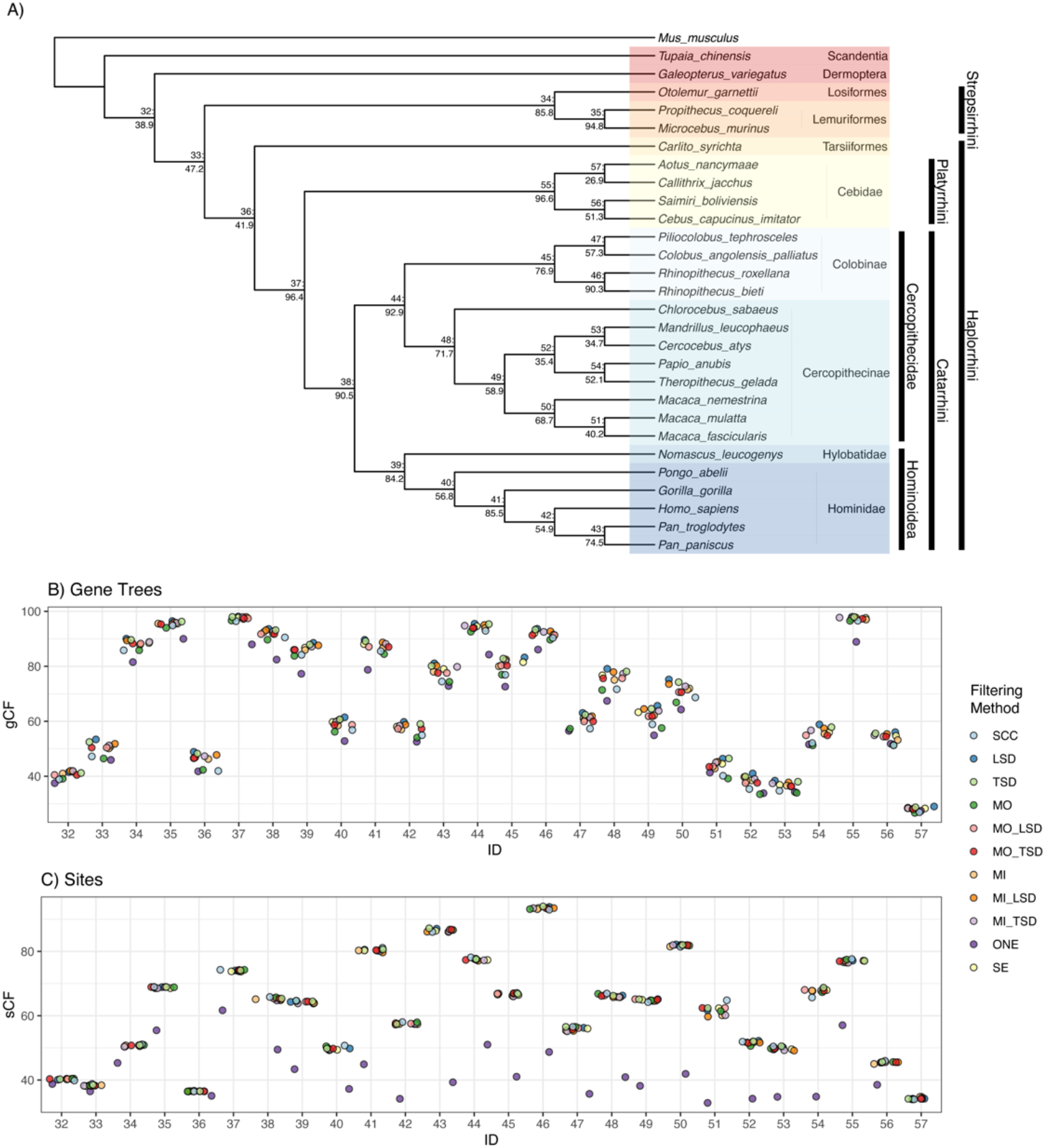
Gene (gCF) and site (sCF) concordance factors among primate datasets using ML gene trees (MIN27). A) Primate phylogeny from ASTRAL-III using the ML gene trees (all input datasets give the same topology). Nodes show Node ID: gCF values from the SCC dataset. B) Distribution of gCF values across datasets. C) Distribution of sCF values across datasets. Node IDs correspond to the numbers displayed on the tree in panel A. SCC=single-copy clusters; LSD=lineage-specific duplicates; TSD=two-species duplicates; MO=monophyletic outgroup; MI=maximum inclusion; SE=subtree extraction; ONE=one paralogs.

### Species tree inference is largely consistent across primate datasets

We inferred species trees using seven approaches: ASTRAL-III (Sayyari and Mirarab 2016; Zhang et al. 2018; Rabiee et al. 2019) on maximum likelihood (ML) gene trees, ASTRAL-III on maximum parsimony (MP) gene trees, ASTRID (Vachaspati and Warnow 2015) on ML gene trees, ASTRID on MP gene trees, concatenated ML inference in IQ-Tree (Nguyen et al. 2015), concatenated MP inference in PAUP* (Swofford 2001), SVDQuartets (Chifman and Kubatko 2014), ASTRAL-Pro (Zhang et al. 2020) on MP and ML gene trees, and ASTRAL-DISCO (Willson et al. 2021) on ML gene trees. ML gene trees were inferred in IQ-Tree, while MP gene trees were inferred in PAUP*. ASTRAL-III, ASTRID, concatenated ML, and concatenated MP were all developed with orthologs in mind, but ASTRAL-III has subsequently been demonstrated to be statistically consistent under models of gene duplication and loss when multiple copies are treated as multiple individuals or when a single copy per species is sampled (Hill et al. 2020; Legried et al. 2020; Markin and Eulenstein 2020). ASTRAL-Pro and ASTRAL-DISCO, on the other hand, were designed with paralogs in mind, and were only applied to the All Paralogs datasets.

Across all nodes of the primate species tree, except for the relationships among the Platyrrhini (discussed below), an identical phylogeny was recovered across all datasets and species tree inference methods (Figure 3), with two exceptions. When concatenated MP or SVDQuartets was used to infer a species tree from the One Paralogs dataset (MIN27), *Macaca fascicularis* was recovered as sister to *M. nemestrina* rather than *M. mulatta*, as in all other datasets and previous studies (e.g., Vanderpool et al. 2020). However, bootstrap support for this relationship was low (55%) in the SVDQuartets analysis. Additionally, when SVDQuartets was used to infer a species tree from the One Paralogs (MIN4) dataset, *Mandrillus leucophaeus* was recovered as sister to a clade containing *Cercocebus atys, Papio anubis*, and *Theropithecus gelada*, rather than sister to *Cercocebus atys* as in other analyses and previous studies; bootstrap support for this relationship was also low (< 50%).

Branch support values were also highly similar across filtering methods. Local posterior probabilities were 1.0 in ASTRAL-III for all datasets and for all nodes except the contentious node in the Platyrrhini. All local posterior probabilities were also 1.0 in ASTRAL-DISCO. All bootstrap support values in the concatenated ML analyses were 100, and all bootstrap support values were 100 in the concatenated MP analyses except for in the One Paralogs (MIN27) dataset, which also had topological issues among macaques as mentioned above. Similarly, in all SVDQuartets analyses bootstrap values were 99 or 100 except among the Platyrrhini and in the One Paralogs datasets.

In addition to branch support values, we calculated measures of genealogical discordance: gene and site concordance factors (gCFs and sCFs; Minh et al. 2020). These analyses were carried out for all datasets except All Paralogs, because it is not possible to calculate these statistics for this dataset in IQ-Tree, which requires a single sample per taxon. For all datasets except the One Paralogs dataset, site and gene concordance factors were highly similar across datasets (Figure 3A, B, C). Concordance in the One Paralogs dataset was consistently lower, as would be expected from the random sampling of homologs. In some cases, gene concordance factors were slightly lower for the SCC and MO dataset than for the other datasets (Figure 3B); this seems to be due to more genes that fall into the ‘paraphyly’ category (i.e., genes for which at least one of the reference clades for a particular branch is not monophyletic), rather than for more genes supporting either of the two minor topologies. Gene and site concordance factors for the MIN4 datasets are shown in Supplemental Figure S2.

### Resolution of the Platyrrhini radiation varies across species tree and gene tree inference methods

As in Vanderpool et al. (2020), we found uncertainty around relationships among the Platyrrhini. Concatenated ML analyses and gene-tree based analyses that relied on gene trees inferred using ML preferred a symmetric tree, with *Saimiri boliviensis* and *Cebus capucinus imitator* as sister species and *Callithrix jacchus and Aotus nancymaae* as sister species (topology 1 in Figure 4A). However, concatenated MP and gene-tree based analyses that relied on gene trees inferred using MP preferred an asymmetric topology, with *S. boliviensis* and *C. c. imitator* sister and *A. nancymaae* sister to these two (topology 2 in Figure 4A). Finally, SVDQuartets preferred a third topology that placed *C. jacchus* sister to *S. boliviensis* and *C. c. imitator* (topology 3 in Figure 4A).

**Figure 4.**
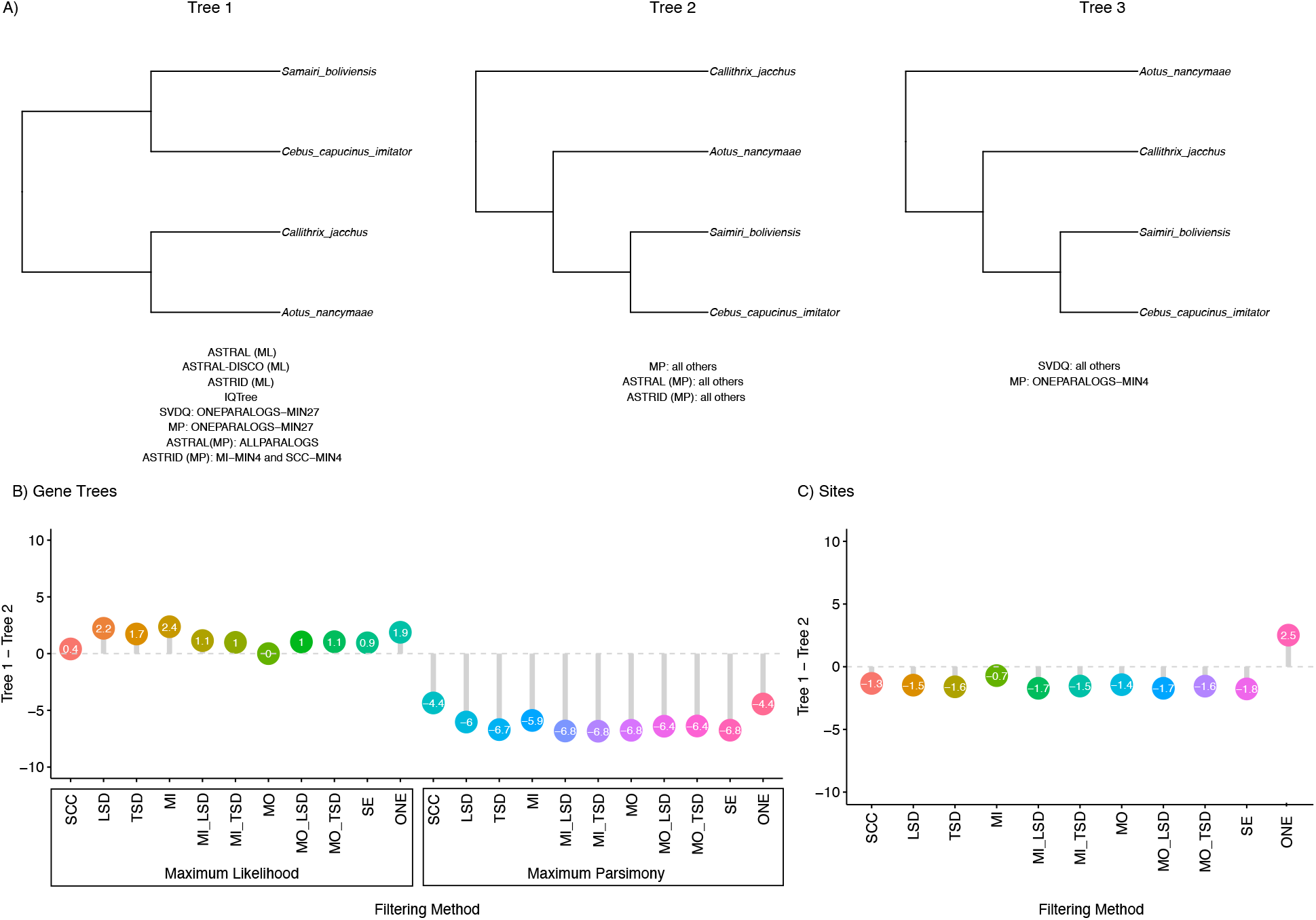
Alternative resolutions of Platyrrhini relationships. A) The three most common tree topologies. Below each resolution, inference methods and filtering approaches that supported the topology are listed. B) The percentage of gene trees supporting Tree 1 minus the percentage of gene trees supporting Tree 2 for ML and MP gene trees across datasets. C) The percentage of sites supporting Tree 1 minus the percentage of sites supporting Tree 2 across datasets. SCC=single-copy clusters; LSD=lineage-specific duplicates; TSD=two-species duplicates; MO=monophyletic outgroup; MI=maximum inclusion; SE=subtree extraction; ONE=one paralogs. Results in B and C from MIN27 datasets.

Gene and site concordance factors clarify these results. A slight majority of ML gene trees prefer topology 1 (Figure 4B), a majority of MP gene trees prefer topology 2 (Figure 4B), while slightly more sites support topology 2 than support topology 1 (Figure 4C). While the results from SVDQuartets may seem counterintuitive at first, SVDQuartets relies on symmetry between the two minor topologies to infer the third topology as the correct topology. Since there are relatively equal numbers of sites supporting topologies 1 and 2, it is therefore expected that SVDQuartets would prefer topology 3, even though fewer sites support this topology. Results for the MIN4 dataset are similar and are shown in Figure S3.

To further investigate the causes of disagreement among these taxa, we focused on the SCC dataset with MIN27 filtering to compare ML and MP gene trees. For each gene, we recorded the ML and MP gene tree topology and the sCF with respect to the focal node, as well as various summary statistics about each locus (number of site patterns, number of parsimony informative sites, tree length, etc.). The percentage of sites supporting the best topology was highest when ML and MP gene trees agreed (Supplemental Figure S6a,c). Additionally, there was more variance in sCFs within a gene (i.e., the number of sites supporting each topology differed more) when ML and MP gene trees agreed (Supplemental Figure S6a,b). This suggests that for genes with similar numbers of sites supporting multiple topologies, ML and MP were more likely to infer conflicting gene trees. Notably, 17.6 percent of gene trees supported Tree 1 under both ML and MP inference, while 18.8 percent of the gene trees supported Tree 2 under both ML and MP inference.

### Branch length estimates are largely consistent across primate datasets

We inferred branch lengths using two approaches. In general, our results suggest that all methods that extract orthologs perform similarly and should lead to reliable estimates of branch lengths. First, we estimated branch lengths in units of substitutions per site using concatenated ML (i.e., site-based branch lengths). We expect that the inclusion of paralogs will lead to overestimates of site-based branch lengths, since the divergence times of paralogs should pre-date divergence times of orthologs. As expected, estimated site-based branch lengths for the One Paralogs dataset are longer than those estimated for the SCC dataset (Figure 5a, b). For all other MIN27 datasets, estimated site-based branch lengths were highly similar to those from the SCC dataset (Figure 5 c, d). However, there are some inconsistencies with site-based branch lengths for terminal branches (Figure 5d), and all site-based branch lengths are more variable for the MIN4 datasets (Supplemental Figure S4).

**Figure 5.**
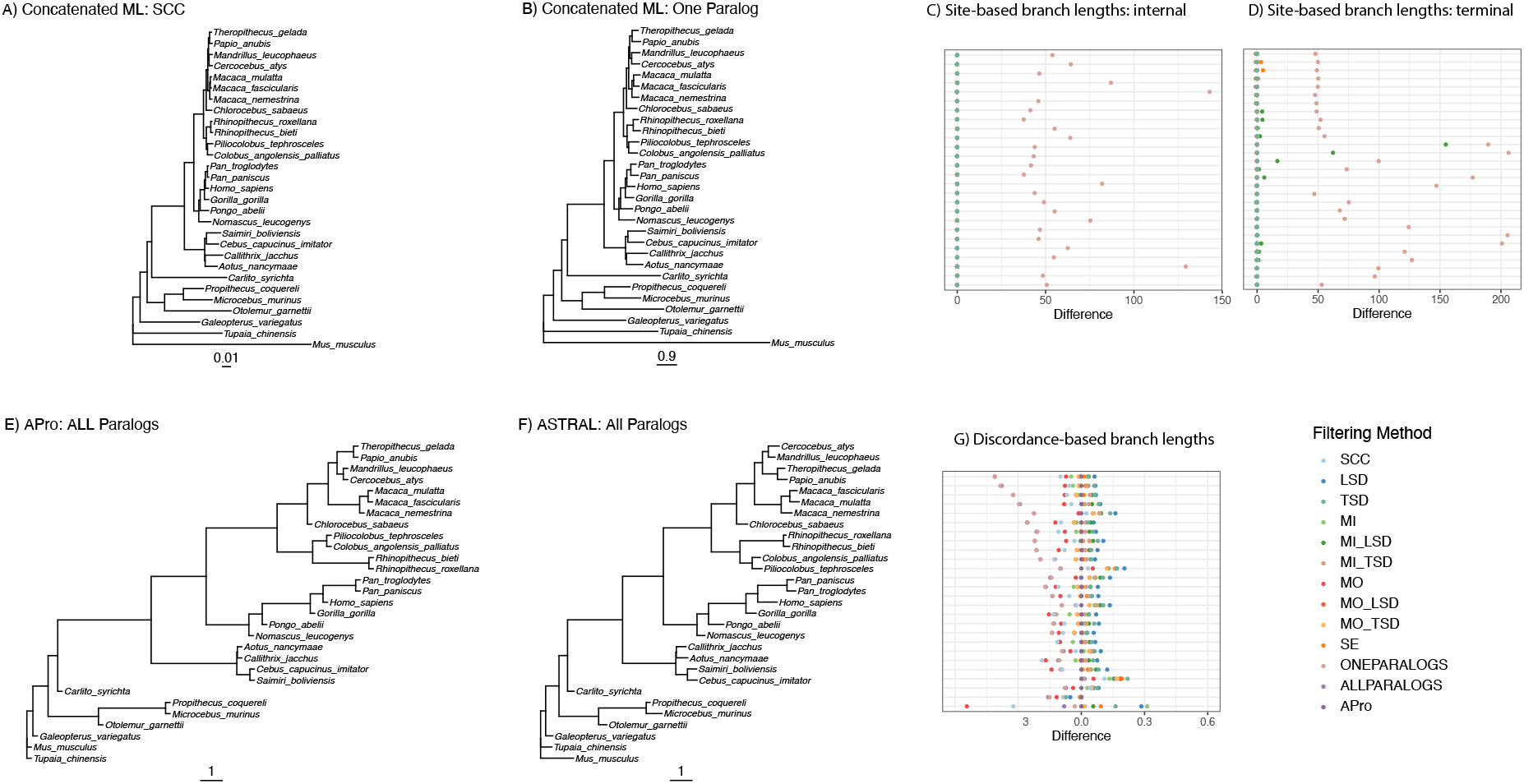
Branch lengths across primate datasets and species tree inference methods. Site-based branch lengths estimated using concatenated ML when A) SCCs and B) one randomly selected paralog per species are used for inference. Note the different scales in panels A and B. C) Difference between site-based branch lengths for internal branches from the SCC dataset and all other datasets, normalized by SCC branch length. D) Same as in panel C, but for terminal branches. Discordance-based branch lengths calculated on the All Paralogs dataset when E) ASTRAL-Pro and F) ASTRAL-III are used for inference. Note that terminal branch lengths are arbitrary in these panels. G) Difference between discordance-based branch lengths estimated with ASTRAL-Pro (APro) and all other methods, normalized by APro branch length. Colors represent different filtering methods, and each row is a different branch. SCC=single-copy clusters; LSD=lineage-specific duplicates; TSD=two-species duplicates; MO=monophyletic outgroup; MI=maximum inclusion; SE=subtree extraction; ONE=one paralogs. Results from MIN27 datasets.

We also inferred discordance-based branch lengths in coalescent units using ASTRAL-III for the ML gene tree datasets. We expect that the inclusion of paralogs will lead to underestimated discordance-based branch lengths, because datasets with paralogs should have higher levels of discordance. As expected, estimated discordance-based branch lengths from the All Paralogs and One Paralogs dataset using ASTRAL-III are shorter than those estimated from the All Paralogs dataset using ASTRAL-Pro, a method that accounts for the extra discordance caused by the inclusion of paralogs (Figure 5 e, f, g). In general, across all datasets except the two including paralogs (All and One), discordance-based branch lengths were highly similar to those estimated in ASTRAL-Pro (Figure 5g). However, there were some surprising results.

Specifically, the SCC and MO datasets led to slightly shorter discordance-based branch length estimates than both ASTRAL-Pro and the datasets from other tree-based decomposition methods (Figure 5g). In addition, all discordance-based branch length estimates are relatively short, which could be explained by difficulties estimating the lengths of longer branches with very little gene tree discordance (i.e., for which all (or most) genes support a single topology) in ASTRAL-III.

### Tests for introgression are consistent across primate datasets

To test for introgression, we looked for a deviation from the expected number of alternate gene tree topologies using the statistic Δ (Huson et al. 2005; Vanderpool et al. 2020). We used only the ML gene trees from each dataset for this analysis. There was evidence of introgression across several branches of the primate phylogeny (Figure 6A), and values of Δ were similar across datasets (Figure 6B). Notably, there was evidence of introgression in a majority of tests at the contentious node in the Platyrrhini, which may explain difficulties inferring the species tree topology at this node. There was also evidence of introgression in the macaques, as found by Vanderpool et al. (2020). Deeper in the tree, results were more suspect, with tests on some datasets suggesting introgression while others did not (Figure 6B). Results of introgression tests were similar with less stringent missing-data filters (Supplemental Figure S5).

**Figure 6.**
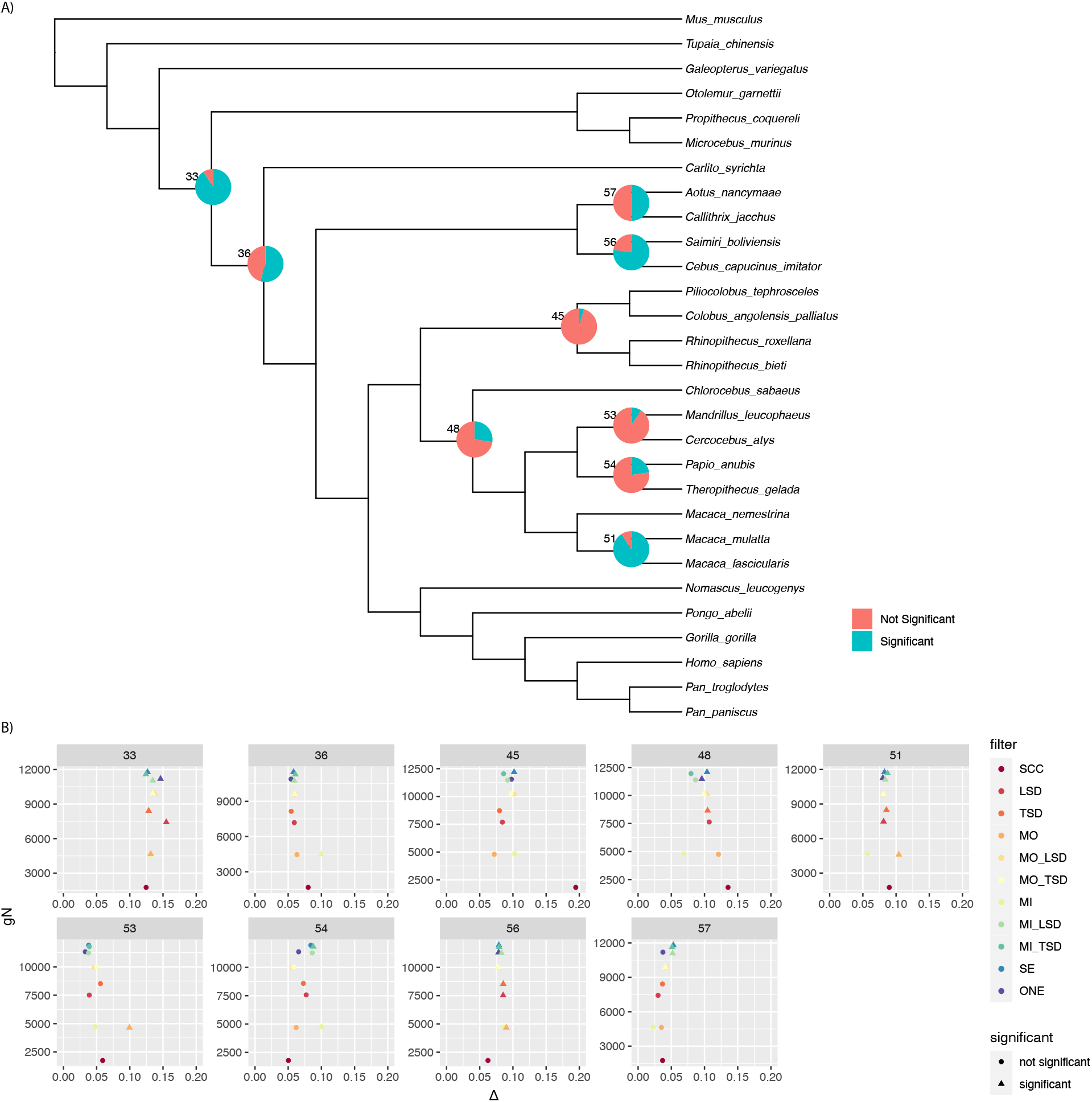
Results of introgression tests on primate MIN27 ML gene trees. A) Pie charts are shown for branches with any significant introgression tests. Numbers are node numbers. B) For all branches with some significant tests, we show the number of informative genes versus Δ. Observations are colored by filtering method, and shapes indicate whether a particular test was significant. SCC=single-copy clusters; LSD=lineage-specific duplicates; TSD=two-species duplicates; MO=monophyletic outgroup; MI=maximum inclusion; SE=subtree extraction; ONE=one paralogs.

### Inferred species trees are largely consistent across additional clades

We assembled datasets and inferred species trees for several other empirical datasets previously analyzed by Morel et al. (2022). We analyzed five datasets: a fungi dataset including 16 species (fungi-16; Rasmussen and Kellis 2012), a fungi dataset including 60 species (fungi-60; Huerta-Cepas et al. 2014), a vertebrate dataset including 22 species (vertebrates-22; Huerta-Cepas et al. 2014), a vertebrate dataset including 188 species (vertebrates-188; Zerbino et al. 2018), and a plant dataset including 23 species (plants-23; Huerta-Cepas et al. 2014). These datasets varied widely in the number of gene copies (Appendix B). The proportion of gene families that were single-copy ranged from ∼3% in the plants-23 dataset to ∼67% in the fungi-16 dataset. The datasets also varied in the number of gene families (Appendix B), the number of taxa, and the depth of divergence. For each dataset, we assembled seven subsets of gene families: single-copy clusters (SCC), lineage-specific duplicates (LSD), two-species duplicates (TSD), MI-extracted orthologs with two-species duplicates removed (MI-TSD), SE-extracted orthologs, All Paralogs, and One Paralogs. We then inferred species trees using ASTRAL-III, ASTRAL-Pro, ASTRID, concatenated ML, and concatenated MP. For three datasets, ASTRAL-III could not complete using the memory and wall-time available (up to 500 Gb and 94 hours), so for these datasets we used a modified version of FASTRAL (Dibaeinia et al. 2021). We omit results from other analyses that did not complete with 94 hours of wall-time and 500 Gb of memory (Appendix B).

In general, across any given inference method (e.g., all trees inferred with ASTRAL-III), species tree topologies were highly similar whether we used single-copy clusters or orthologs extracted from larger gene families (Figure 7; Appendix B). The largest differences were between trees inferred using concatenated ML and concatenated MP on the one hand and those inferred using the gene-tree based methods ASTRAL-III and ASTRID on the other (Appendix B). Analyses of the One Paralogs subset using concatenated approaches resulted in highly different trees for the vertebrates-22 and plants-23 datasets (Appendix B, Figure B4, B7). In two cases, analyzing All Paralogs in ASTRAL-III resulted in different topologies as well. For the fungi-16 dataset, the tree inferred in ASTRAL-III from All Paralogs differed from other trees at contentious nodes, but agreed with some previous studies (Rasmussen and Kellis 2012); nodal support values were also low at these nodes (Appendix B, Figure B1). For the fungi-60 dataset, the tree inferred from All Paralogs using ASTRAL-III was substantially different from other trees; our results suggest that this difference arose due to an issue when searching tree space in ASTRAL-III, rather than due to some inherent property of the dataset (Appendix B). Overall, our results highlight the robustness of topological inference to extracting genes from larger gene families, and, in most cases, to using all data from all gene families.

**Figure 7:**
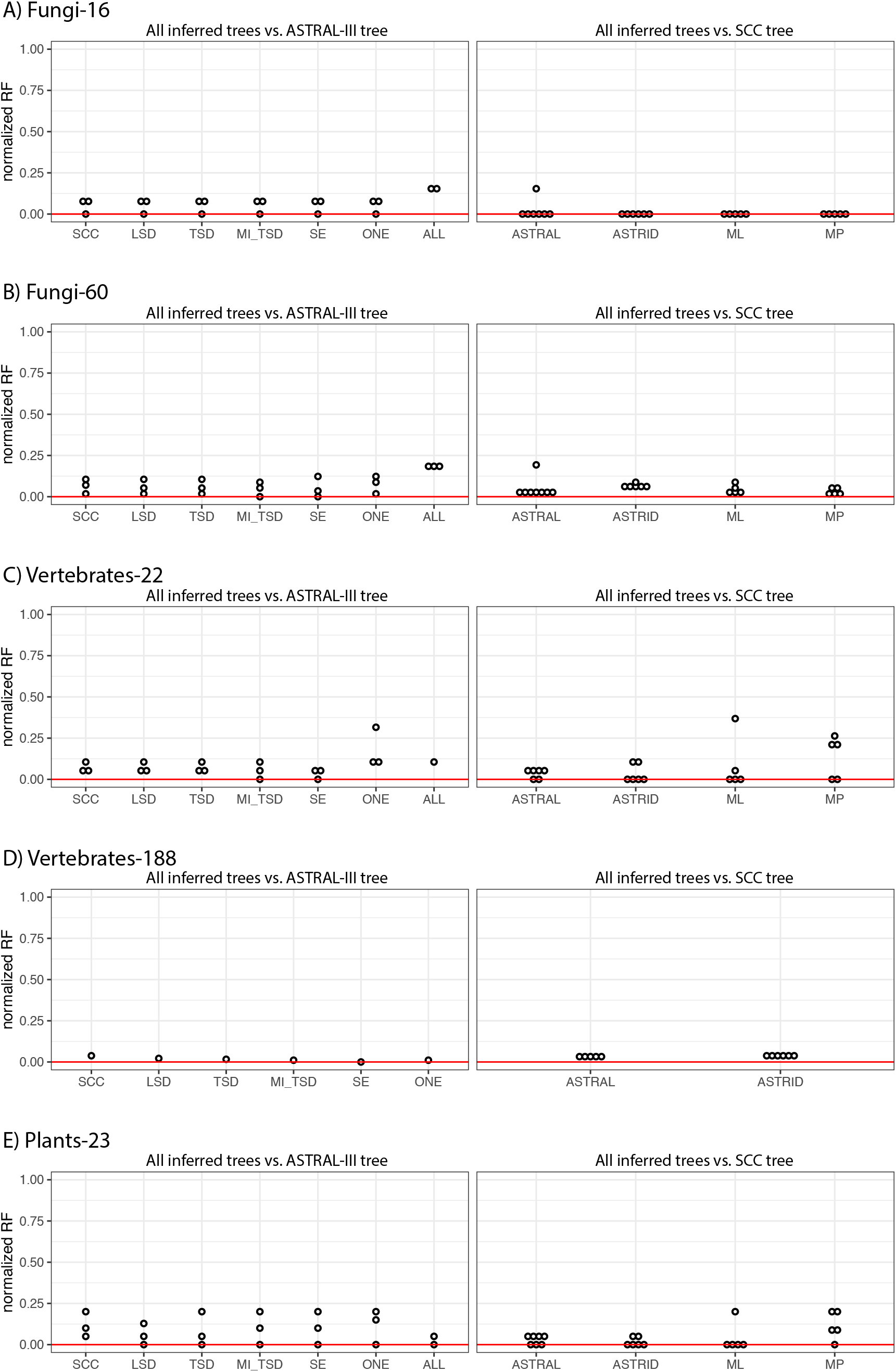
Results from analyzing five additional clades. On the left, we show the normalized Robinson-Foulds distances between trees inferred using different species tree inference methods (ASTRID, concatenated ML, concatenated MP, ASTRAL-Pro) and tree inferred using ASTRAL-III for each data subset. On the right, we show the normalized Robinson-Foulds distances between trees inferred from different data subsets (SCC, LSD, TSD, MI-TSD, SE, ONE, ALL) and the SCC tree for each species tree inference method. In the case of the vertebrates-22 ‘All Paralogs’ data subset, the reference species tree on the left is the ASTRID tree, since ASTRAL-III did not complete. SCC=single-copy clusters; LSD=lineage-specific duplicates; TSD=two-species duplicates; MO=monophyletic outgroup; MI=maximum inclusion; SE=subtree extraction; ONE=one paralogs.

## Discussion

Our results demonstrate that, no matter the subset of the data used, the inferred species tree topology is largely stable; this was especially obvious in our analysis of primate genomes. Regardless of whether all families, families with only a single copy per species, or large families from which orthologs were extracted were used, the only disagreements between trees in the primate analyses were with respect to relationships among the Platyrrhini; in this case the species tree inference method was a larger determinant of results than the particular dataset (Figure 4).

Despite the overall similarity among results, when a single gene was randomly sampled per species results were unstable in two cases, suggesting—unsurprisingly—that such a sampling strategy is not ideal. Among additional datasets sampled from across the eukaryotes, results were also highly consistent whether single-copy clusters or orthologs extracted from larger gene families were used for inference. While using all gene families resulted in consistent estimates of species tree topologies in most cases, analyzing these gene families with methods that were not designed for multi-copy gene families (specifically, ASTRAL-III) resulted in an anomalous result in one case, likely due to issues appropriately searching tree space (Appendix B). Based on the results presented here, when whole genome sequence data are available, using all of the families output by clustering methods followed by the application of gene-tree decomposition methods can greatly expand the data available without sacrificing the accuracy of inference.

Several recent simulation studies have evaluated the impacts of gene duplication and loss on inferences of species tree topologies (Yan et al. 2021; Legried et al. 2020; Zhang et al. 2019; Morel et al. 2022). In studies considering the application of ASTRAL-III to multi-copy gene families (i.e. using ASTRAL-multi), its performance has been surprisingly good given that this method was not designed with duplication and loss in mind (Yan et al. 2021; Legried et al. 2020; Zhang et al. 2019). However, in some cases this approach has been outperformed by methods that explicitly accommodate duplication and loss (Zhang et al. 2019, Willson et al. 2021), likely because these approaches use the information contained within gene duplication events while limiting the effects of noise. ASTRAL-Pro (Zhang et al. 2020) includes an internal reconciliation step that labels speciation and duplication nodes, and is therefore operating similarly to gene tree decomposition approaches that try to identify such nodes in order to extract orthologs (although often not under any explicit model). In a comparison between ASTRAL-Pro and ASTRAL-DISCO (an approach that decomposes gene families prior to analyzing them in ASTRAL-III), ASTRAL-DISCO performed similarly to ASTRAL-Pro with lower computation times (Willson et al. 2021). Similarly, our analyses of six empirical datasets highlight the fact that tree-decomposition approaches perform similarly to ASTRAL-Pro when inferring species tree topologies. Taken together, these results suggest that decomposition is a promising approach for using a wider array of methods to infer species trees from large gene families.

Despite the stability of inference across most of the tree in the primate dataset, there remains disagreement about relationships amongst the Platyrrhini, a notably contentious node (Perelman et al. 2011; Springer et al. 2012; Perez et al. 2013; Jameson Kiesling et al. 2015; Schrago and Seuánez 2019; Wang et al. 2019; Vanderpool et al. 2020). As in Vanderpool et al. (2020), we find that both concatenated ML and ASTRAL-III based on ML gene trees favor a symmetrical topology (tree 1 in Figure 4A). A bias towards the symmetrical 4-taxon tree is expected when using ML in the presence of recombination and when the time between speciation events is short (Kubatko and Degnan 2007; Roch and Steel 2015). Although the bias in ML under these conditions is often linked to concatenation methods, if the gene trees themselves are inaccurate due to the concatenation of multiple unique histories (e.g., among exons; Mendes et al. 2019), then the same bias in inferred trees can occur. Bias in the gene trees can then lead to bias in the methods that they are used as input to (e.g., ASTRAL-III). Note that this bias does not affect inferences under maximum parsimony (Mendes and Hahn 2018). Furthermore, there are nearly equal numbers of trees supporting the two best-supported topologies in the primate data (Figure 4B), which suggests two things: first, choosing the best topology will be difficult no matter what method is used, as the evidence in favor of one topology over the other is minimal. Second, there is likely some introgression, since we would otherwise expect equal numbers of the two minor topologies. We do not see equal numbers of the two minor topologies, as confirmed by significant tests for introgression in this clade (Figure 6). Finally, a detailed comparison of SCC gene trees inferred by both ML and MP suggests that genes whose topologies disagreed across the two approaches did not support either topology as strongly as genes for which ML and MP agreed (Figure S6). Of the gene trees that agreed across ML and MP inference, more supported Tree 2 than supported Tree 1 (Figure 4A). Thus, of the genes for which methods agree, more support the asymmetric topology than the symmetric topology (as in Vanderpool et al. 2020).

We also compared branch length estimates and tests for introgression across datasets. Branch length estimates are largely consistent across datasets, with the exception of datasets that explicitly include paralogs, which led to biases in expected directions for both discordance-based and site-based branch lengths. Site-based branch lengths are very consistent across all datasets except the One Paralogs dataset when stringent filters for missing data are applied. When paralogs are included, site-based branch lengths are overestimated, as expected (e.g., Siu-Ting et al. 2019). Discordance-based branch lengths (i.e., those estimated in ASTRAL) are underestimated for datasets including paralogs, because these datasets have higher levels of discordance. These methods accommodate increased discordance by positing a shorter time between speciation events. Otherwise, discordance-based branch lengths are largely similar across datasets, though the SCC and MO datasets appear to have slightly shorter estimated branch lengths than all other methods (Figure 5e). Given the consistency of results across tree-based decomposition methods, as well as ASTRAL-Pro, and the vastly larger number of gene trees used in these cases, we suggest that discordance-based branch lengths may actually be underestimated for the SCC and MO datasets. This result is consistent with lower gCFs in these datasets (Figure 3b), and suggests that branch lengths estimated from these datasets may be inaccurate because they include pseudoorthologs.

To our knowledge, this is the first evaluation of the effects of including more than just single-copy families on tests for introgression based on the asymmetry in minor topology frequencies. We expected that the inclusion of paralogs would not bias such tests, because under models that include duplication and loss the two minor topologies should occur in equal frequencies (Smith and Hahn 2021a; Smith and Hahn 2021b). Our results largely confirm these expectations: although there is variation in whether or not tests are significant across datasets, estimates of Δ are very similar (Figure 6B). At some nodes there is consistent evidence for introgression across datasets, suggesting a strong signal of asymmetry: for example in the macaques and among the Platyrrhini. Deeper in the tree, there may be more gene tree error (e.g., due to long-branch attraction), since introgression is detected for some datasets and not for others (Figure 6B).

Phylogenetics based on whole-genome sequences almost always begins by identifying homologous genes via clustering. The clustering process operationally defines gene families, using clustering methods that range from very simple to very complex. While the single-copy clusters output by any one of these methods have most often been used in phylogenetics, there is nothing inherently more suitable about these clusters. First, SCCs may not be orthologs, due to the presence of pseudoorthologs—paralogs that are mistaken as orthologs due to differential patterns of gene duplication and loss (Doolittle and Brown 1994; Koonin 2005). In other words, having only a single representative sequence in each species does not guarantee that all the sampled genes are orthologs. Second, and more importantly, the size of clusters identified by clustering approaches is determined by parameters set by the user. For example, in OrthoMCL (Li 2003) the inflation parameter determines the size of output clusters: by changing this parameter, users can identify larger or smaller clusters. Because genes are related to all other genes via a long history of duplication and divergence (with a few exceptions; Knowles and McLysaght 2009; Zhao et al. 2014), there is no single level of similarity that uniquely identifies gene families (Demuth and Hahn 2009). However, users can choose the value of the inflation parameter that identifies more, smaller clusters in order to find more single-copy clusters; this does not mean these genes do not have paralogs, only that more distant paralogs were not included at this clustering threshold. Many clustering methods aim to form groups of genes that descend from a single common ancestor in the studied taxa (e.g., Emms and Kelly 2015), though this does not ensure a lack of duplication events since the common ancestor. While tree-based decomposition approaches still rely on the clustering step to initially identify the homologs from which gene trees are built, their output is directly related to the definitions of orthologs and paralogs, and is more easily interpreted in a phylogenetic context. By applying these decomposition approaches to larger clusters, researchers can avoid arbitrary determinants of which clusters are single-copy and can instead attempt to extract as many sets of orthologs as possible. Not only does this approach increase the amount of data available, but it also uses criteria more directly linked to the evolutionary history of gene families.

Our analyses included genomic datasets across vertebrates, plants, and fungi. While these datasets varied in the number of species, the depth of divergence, and the total number of available gene families, they are all relatively high-quality genomic datasets. Future work should investigate the effects of the inclusion of paralogs using datasets more prone to errors in homology inference and alignment. For example, when transcriptomic data are analyzed, not all homologs will necessarily be sequenced in all species, complicating the identification of orthologs and paralogs, even using tree-based decomposition approaches (Cheon et al. 2020).

Target enrichment-based approaches (e.g., Faircloth et al. 2012; Weitemier et al. 2014) use probes to target specific genomic regions and may inadvertently capture paralogous sequences. These data are generally limited to a moderate number of targeted orthologous regions, and the incidental inclusion of paralogs may have a much more pronounced effect, as there is far less signal available to overcome the noise associated with incorrect inferences of homology. Finally, inferences of homology may be more difficult when deeper phylogenetic problems are considered and in groups with frequent allo- and autopolyploidy. These scenarios may challenge current phylogenomic methods in ways that the genomic datasets analyzed here do not, and should be carefully considered in future work.

In conclusion, our results suggest that methods for species tree inference are accurate across datasets, whether single-copy clusters or tree-based decomposition methods are used. For most subsets of the data and inference methods, using all clusters (i.e. paralogs and orthologs) also results in consistent inferences of species tree topologies. Our results highlight the benefits of using data from all gene families by showing that the amount of data used can be increased by an order of magnitude (Table 1; Figure 2; Appendix B). While even the smallest dataset was sufficient for accurate species tree inference in the datasets analyzed here, that is not always the case (e.g., Emms and Kelly 2018; Thomas et al. 2020). In such cases, using only single-copy clusters may be not possible, and using data from larger gene families will be essential. Finally, more data facilitates inferences beyond species tree topology, including branch length estimates and the detection of introgression. Our results suggest that branch lengths estimated from single-copy clusters may be less consistent than those estimated using data from larger gene families in the primate dataset (Figure 5), and adding gene families improves our ability to detect significant deviations from symmetric minor topology counts in tests for introgression (Figure 6). Our results are consistent across six empirical datasets that differ in the number of species, the number of gene families, the sizes of gene families, and the depth of divergence. While these datasets are not exhaustive, they suggest the potentially broad applicability of our findings, particularly with respect to the suitability of orthologs extracted from larger gene families for inferring species tree topologies.

## Materials and Methods

### Primate dataset and alignment

The full sets of protein-encoding genes for 26 primates and 3 non-primates were obtained as in (Vanderpool et al. 2020), and clusters were obtained as in that study. Briefly, an all-by-all BLASTP search (Altschul et al. 1990; Camacho et al. 2009) was executed, and the longest isoform of each protein-coding gene from each species was used. Then, the mcl algorithm (Van Dongen 2000) as implemented in FastOrtho (Wattam et al. 2014) with an inflation parameter of 5 was used to cluster the BLASTP output. CDSs for each cluster that included samples from at least four species were aligned, cleaned and trimmed as in (Vanderpool et al. 2020). Sequences were aligned by codon using GUIDANCE2 (Sela et al. 2015) with MAFFT v7.407 (Katoh and Standley 2013) with 60 bootstrap replicates. Sequence residues with GUIDANCE scores < 0.93 were converted to gaps and sites with >50% gaps were removed using Trimalv1.4rev22 (Capella-Gutiérrez et al. 2009). GUIDANCE2 uses the command ‘mafft --localpair --maxiterate 1000 --nuc –quiet’ when running MAFFT. Alignments shorter than 200 bp and alignments that were invariant or contained no parsimony informative characters were removed from further analyses. A subset of alignments that could not be aligned by codon were aligned by nucleotide, and subsequent steps were as with the codon-aligned dataset. In total 18,484 alignments were used in downstream analyses.

### Gene tree inference

We inferred gene trees from all alignments with at least four species (18,484 alignments) in IQ-TREE v2.0.6 (Nguyen et al. 2015) with nucleotide substitution models selected using ModelFinder (Kalyaanamoorthy et al. 2017) as implemented in IQ-TREE. The full IQ-TREE command used on each alignment was ‘iqtree2 -s *alignment name* -m MFP -c 1 -pre *alignment name*’. We also inferred gene trees from all 18,484 alignments using the maximum parsimony criterion in PAUP* v 4.0a (Swofford 2001). We treated gaps as missing data, obtained a starting tree via random stepwise addition, held a single tree at each step, and used the TBR branch-swapping algorithm with a reconnection limit of 8. We kept a maximum of 1000 trees and did not collapse zero-length branches.

### Filtering

We considered three major groups of filtering methods:

1. Single-copy clusters (SCCs): We considered a dataset that consisted only of those clusters that included a single gene copy from each species.
2. Tree-based decomposition approaches: We considered several methods that involved trimming branches of gene trees to extract orthologs. All custom branch-cutting operations were written in python3 and used the python package ete3 (Huerta-Cepas et al. 2016) to read, traverse, trim, and output gene trees and modified sequence alignments. We used postorder node traversal when traversing trees, and prior to custom trimming operations, we midpoint-rooted gene trees.
  i. Lineage-specific duplicates: In this dataset we identified gene duplications that were specific to a single species. For such lineage-specific duplicates, we selected the sequence copy that was closest in length to the median length of sequences in the alignment, kept that copy, and trimmed the other copy or copies from both the alignment and the gene tree.
  ii. Two-species duplicates: To expand our data beyond lineage-specific duplicates, in addition to trimming lineage-specific duplicates, we identified gene duplications specific to a pair of species. For such duplicates, we selected the two sequence copies with the minimum branch distance separating them and trimmed the remaining copies from the tree and the alignment.
  iii. Maximum Inclusion: We applied the maximum inclusion (MI) approach described in (Yang and Smith 2014) to trim gene trees. We used the python script provided by Yang and Smith (2014; prune_paralogs_MI.py) and used as input one of three sets of gene trees: the original 18,484 gene trees, the original 18,484 gene trees with lineage-specific duplicates trimmed, and the original 18,484 gene trees with lineage-specific and two-species duplicates trimmed. For the MI approach, branches longer than a specified threshold are trimmed to remove potential pseudoorthologs; we used the following branch-length cutoffs: 0.4 substitutions per site for the ML gene trees and 500 changes for MP trees. We explored additional cutoffs in the Supplementary Material (Appendix A).
  iv. Monophyletic Outgroups: We also applied the monophyletic outgroups (MO) approach described in (Yang and Smith 2014) to trim gene trees. We used the python script provided by Yang and Smith (2014; prune_paralogs_MO.py) and used as input one of three sets of gene trees: the original 18,484 gene trees, the original 18,484 gene trees with lineage-specific duplicates trimmed, and the original 18,484 gene trees with lineage-specific and two-species specific duplicates trimmed.
  v. Subtree Extraction: Finally, we evaluated a new tree-based decomposition approach introduced here (subtree extraction; SE). In this approach we start by midpoint-rooting gene trees, followed by trimming lineage-specific and two-species duplicates. We then extract subtrees with a single representative from each taxon (i.e., subtrees with no duplicates) and keep those subtrees that meet minimum taxon-sampling thresholds.
3. Paralog methods: We considered two approaches that included paralogs in addition to orthologs. First, we included all genes (All Paralogs). Additionally, we randomly sampled a single gene (without regard to orthology) per species (One Paralogs).

For all datasets, we considered a stringent (minimum of 27 of 29 taxa) and relaxed (minimum of 4 of 29 taxa) missing data threshold.

### Species tree inference

We inferred species trees using seven methods. Three methods inferred species trees from concatenated datasets: maximum parsimony, maximum likelihood, and SVDQuartets. To infer an MP tree from the concatenated datasets we used PAUP* v4.0a (build 168) (Swofford 2001). We set the criterion to parsimony, and used 500 bootstrap replicates to assess nodal support. For all other options we used PAUP* defaults. To infer an ML tree from the concatenated dataset we used IQ-TREE v2.0.6 (Nguyen et al. 2015) with nucleotide substitution models selected using ModelFinder (Kalyaanamoorthy et al. 2017) as implemented in IQ-TREE. We used an edge-linked, proportional partition model (Chernomor et al. 2016) and 1000 ultrafast bootstrap replicates (Hoang et al. 2018). The full IQ-TREE command used on each alignment was ‘iqtree2 -s *alignment name* -p *partition file name* -c 1 -pre *alignment name -B 1000*’. For three alignments, IQTree v2.0.6 failed to run, and, based on a suggestion from the developers, we reverted to IQTree v.1.6.12 to infer the species trees for these alignments. For these three alignments, the full IQ-TREE command used was ‘iqtree -s *alignment name* -spp *partition file name* -pre *alignment name* -bb 1000 -nt 4’. Finally, to infer a species tree from the concatenated alignments using SVDQuartets, we used PAUP* v4.0a (build 168) (Swofford 2001). We evaluated all quartets and treated ambiguous sites as missing to infer the species tree topology using the command ‘svdq evalq=all bootstrap=no ambigs=missing loci=allchars;’. To assess nodal support, we evaluated 10,000 random quartets for each of 100 bootstrap replicates. We used the multilocus bootstrapping option and again treated ambiguous sites as missing. The command used for bootstrapping in SVDQuartets was ‘svdq evalq=random nquartets=10000 bootstrap=multilocus loci=allchars nreps=100 nthreads=2 replace=yes treefile=*output file name* ambigs=missing;’.

In addition to the three concatenation-based methods, we inferred species trees using four gene-tree based methods. Prior to inferring species trees or estimating discordance (see below) from filtered gene trees, we collapsed all zero-length branches. For each gene tree, we did the following: First, we midpoint-rooted the gene tree. Then, we calculated sCFs using IQ-Tree v2.0.6 (Minh et al. 2020) for the alignment with the rooted gene tree as the reference tree. We used 100 randomly sampled quartets to compute the sCF, collapsing any nodes where sN == 0; in other words, any nodes for which no sites were informative.

We inferred a species tree using ASTRAL-III v5.7.3 (Sayyari and Mirarab 2016; Zhang et al. 2018; Rabiee et al. 2019). ASTRAL-III infers a species tree from a set of gene trees by extracting quartets and finding the species tree that maximizes the number of shared quartet trees. It has been demonstrated to be consistent under the multispecies coalescent (MSC) model (Mirarab et al. 2014) and under models of gene duplication and loss (Legried et al. 2020). Gene trees obtained using ML and MP, from all datasets described above, and with zero-length branches collapsed, were used as input to ASTRAL-III; local posterior probabilities were used to assess nodal support. In order to run ASTRAL-III on mutli-copy gene trees (i.e. the All Paralogs dataset), we used the mapping file and treated each gene copy as a separate individual.

Additionally, we inferred species trees using ASTRID v2.2.1 (Vachaspati and Warnow 2015), again using the filtered and zero-length collapsed ML and MP gene trees as input. ASTRID is a distance-based approach that estimates species trees using internode distances and is statistically consistent under the MSC model (Vachaspati and Warnow 2015). As in ASTRAL-III, for the All Paralogs dataset we treated gene copies from the same species as individuals using the mapping file. Finally, we inferred species trees from the All Paralogs datasets using ASTRAL-Pro (Zhang et al. 2020) and ASTRAL-DISCO (Willson et al. 2021). ASTRAL-Pro uses an internal rooting-and-tagging algorithm to label nodes as duplication or speciation nodes, and then infers quartets using only speciation nodes before finding the species tree that maximizes the number of shared quartet trees. ASTRAL-Pro has been shown to be statistically consistent under a model of gene duplication and loss, provided that rooting and tagging of nodes as speciation or duplication nodes is correct (Zhang et al. 2020). ASTRAL-DISCO decomposes multi-copy gene trees into single-copy trees using the “rooting and tagging” algorithm from ASTRAL-Pro and then infers a species tree using ASTRAL-III.

### Assessing discordance

To assess levels of discordance across datasets we calculated gene and site concordance factors in IQ-Tree v2.0.6 (Minh et al. 2020). We used as the reference tree the tree shown in Figure 3, and to estimate sCFs, we used 1000 randomly sampled quartets. gCFs were estimated for filtered ML and MP gene trees after zero-length branches were collapsed. sCFs were estimated for the alignments that resulted from filtering the ML gene trees.

### Testing for introgression

We used the approach used in (Vanderpool et al. 2020) to test for introgression. Briefly, the introgression test assesses whether there is a deviation from the expected equal numbers of alternative tree topologies (under the MSC model without gene flow) using the statistic Δ (Huson et al. 2005), where

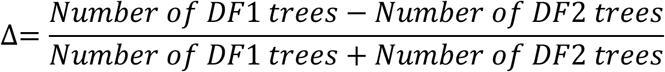

*DF1* represents the most common minor topology, and *DF2* represents the least common minor topology. In the absence of introgression, Δ is expected to be equal to zero. To test whether deviations from zero were significant, we followed the procedure of (Vanderpool et al. 2020) and used 2,000 datasets generated by resampling gene trees with replacement, considering only those nodes where more than five percent of trees were discordant. This distribution was used to calculate *Z*-scores and *p*-values for the observed Δ statistic, and for each filtered dataset, we corrected for multiple comparisons using the Dunn-Sidák correction (Dunn 1959; Šidák 1967).

### Fungi, vertebrate, and plant datasets

We downloaded the fungi-60, vertebrates-22, vertebrates-188, and plants-23 datasets from Morel et al. (2022). The fungi-60, vertebrates-22, and plants-23 datasets were extracted from the PhlomeDB database (Huerta-Cepas et al. 2014) by Morel et al. (2022). For these three datasets, amino acid matrices were used in concatenated analyses. We used gene trees from Morel et al. (2022) inferred from amino acid matrices using ParGenes (Morel et al. 2019) and RAxML-NG (Kozlov et al. 2019) for the fungi-60 and plants-23 datasets. For the vertebrates-22 dataset, we followed Morel et al. (2022) in using the gene trees from the PhylomeDB database, which were reconstructed in PhyML v3.0 (Guindon and Gascuel 2003) from amino aid matrices. The vertebrates-188 dataset was extracted from the Ensembl Compara database (Zerbino et al. 2018) by Morel et al. (2022). For this dataset, nucleic acid matrices were used for concatenated analyses. We used gene trees from Morel et al. (2022) inferred from nucleic acid matrices using ParGenes (Morel et al. 2019) and RAxML-NG (Kozlov et al. 2019). We downloaded the fungi-16 dataset (Rasmussen and Kellis 2012) from http://compbio.mit.edu/dlcoal/. For this dataset, nucleic acid alignments were used for concatenated analyses, and we used gene trees from the original study inferred from nucleic acid matrices using PhyML (Guindon and Gascuel 2003). We removed two trees that had polytomies.

For each dataset, we assembled seven subsets of gene families: single-copy clusters (SCC), lineage-specific duplicates (LSD), two-species duplicates (TSD), MI-extracted orthologs with two-species duplicates removed (MI-TSD), SE-extracted orthologs, All Paralogs, and One Paralogs. We inferred species trees using ASTRAL-III (Sayyari and Mirarab 2016; Zhang et al. 2018; Rabiee et al. 2019), ASTRID (Vachaspati and Warnow 2015), ASTRAL-Pro (Zhang et al. 2020), concatenated MP inference in PAUP* (Swofford 2001), and concatenated ML Inference in IQ-Tree (Nguyen et al. 2015). When ASTRAL-III could not complete with 94 hours and 500 Gb, we ran FASTRAL (Dibaeinia et al. 2021). In order to run FASTRAL on datasets with missing data, we made slight changes to the FASTRAL source code by automating the construction of a custom map file for each run of ASTRID. We calculated distances between inferred trees using the python package ete3 (Huerta-Cepas et al. 2016).

## Acknowledgements

This work was supported by a National Science Foundation postdoctoral fellowship to MLS (DBI-2009989) and an NSF grant to MWH (DEB-1936187).

## Data Availability

Scripts used for filtering gene trees are available on GitHub (github.com/meganlsmith/Primate_Paralogs). Primate alignments, gene trees, and species trees are available from FigShare (doi: 10.6084/m9.figshare.16653025).

