## Supporting Information for "Using all gene families vastly expands data available for phylogenomic inference"

**Includes:**

Supporting Figures S1-S5

Appendix A: Alternative MI datasets (Table A1, Figures A1-A7)

Appendix B: Analyses of non-primate datasets (Tables B1-B5; Figures B1-B8)

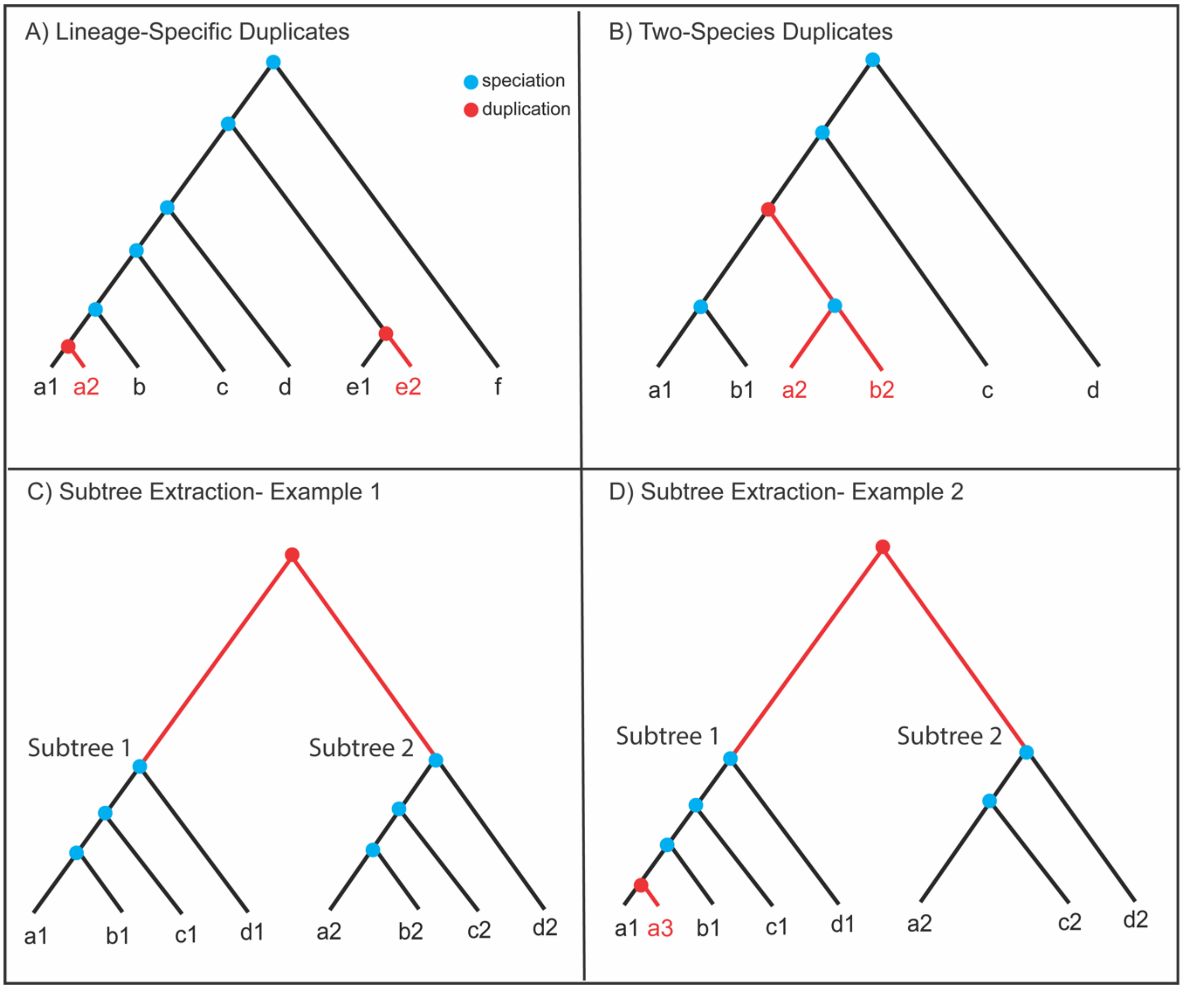


**Figure S1**. Examples of tree-based decomposition approaches. Clipped branches are drawn in red, and sequences omitted from the final dataset are labeled in red. A) Lineage-Specific Duplicates (LSDs). When a duplication occurred in the ancestor of a single species, we can select one copy to keep for downstream analyses (a1, e1) and remove the other copy (a2, e2). All remaining copies are orthologs. B) Two-Species Duplicates (TSDs). When a duplication occurred in the ancestor of two species, we can select one of the two duplicated subtrees to keep for downstream analyses, and remove the other copies (a2 and b2). In the example shown only orthologs remain. C) Subtree Extraction (SE). This approach extracts subtrees that do not contain taxon duplicates. In this case, we extract two trees, one with a1, b1, c1, and d1, and the other with a2, b2, c2, and d2. D) Subtree extraction automatically clips LSDs and TSDs, as shown in subtree 1, when copy a3 is removed. Subtree extraction also works in the case where some taxa are missing, as illustrated in subtree 2, where there is no copy in taxon B.

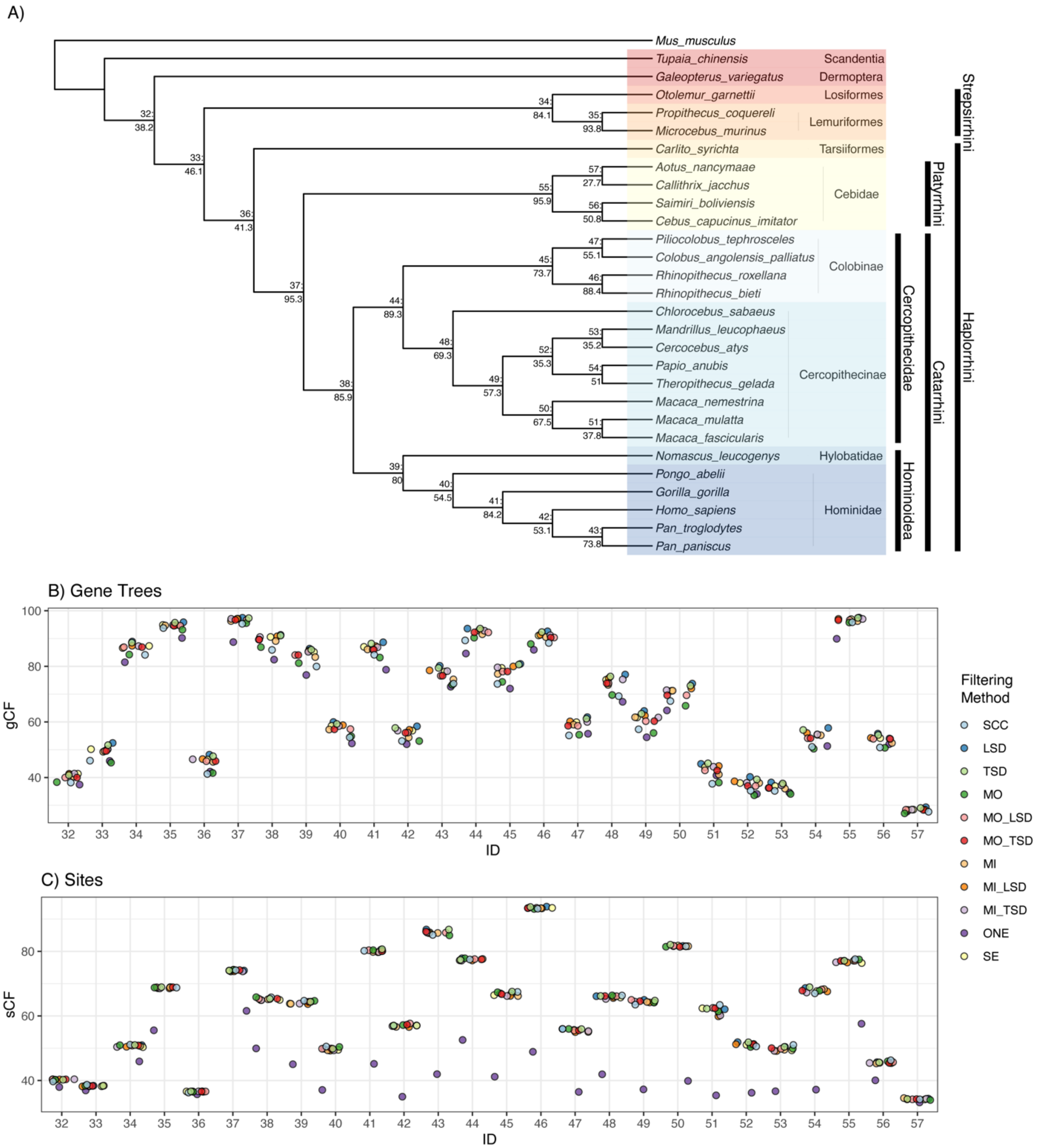


**Figure S2**. Gene (gCF) and site (sCF) concordance factors among primate datasets using ML gene trees (MIN4). A) Primate phylogeny from ASTRAL-III using the ML gene trees (all input datasets give the same topology). Nodes show Node ID: gCF values from the SCC dataset. B) Distribution of gCF values across datasets. C) Distribution of sCF values across datasets. Node IDs correspond to the numbers displayed on the tree in panel A. SCC=single-copy clusters; LSD=lineage-specific duplicates; TSD=two-species duplicates; MO=monophyletic outgroup; MI=maximum inclusion; SE=subtree extraction; ONE=one paralogs.

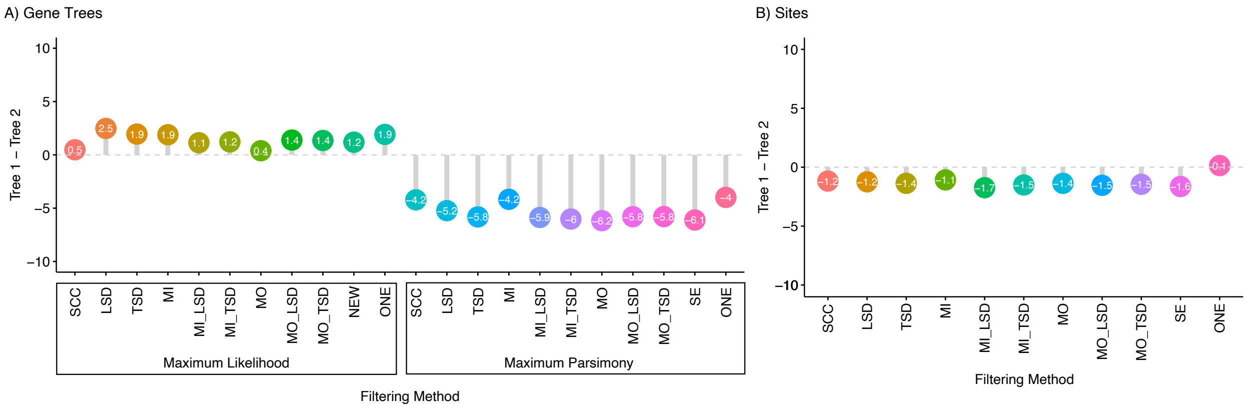


**Figure S3.** Alternative support for Platyrrhini relationships (MIN4 dataset). A) The percentage of gene trees supporting Tree 1 minus the percentage of gene trees supporting Tree 2 for ML and MP gene trees across datasets (refer to Figure 4 in the main text for the tree topologies). B) The percentage of sites supporting Tree 1 minus the percentage of sites supporting Tree 2 across datasets. SCC=single-copy clusters; LSD=lineage-specific duplicates; TSD=two-species duplicates; MO=monophyletic outgroup; MI=maximum inclusion; SE=subtree extraction; ONE=one paralogs.

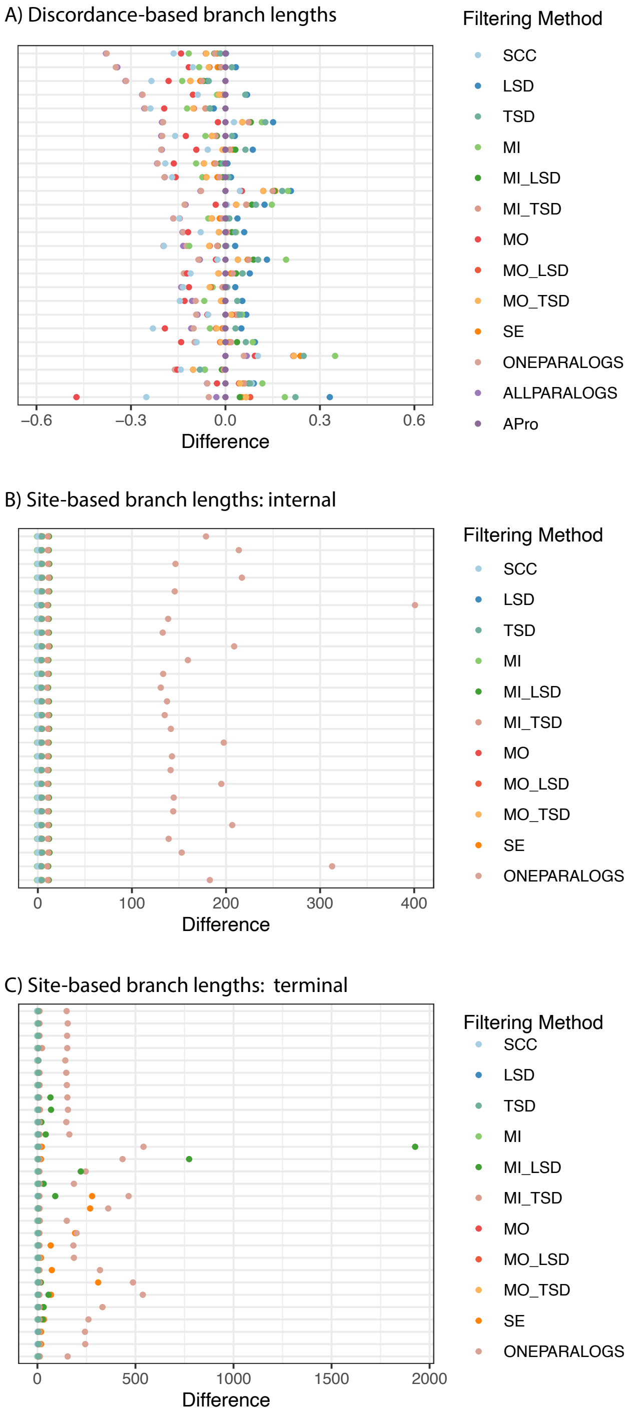


**Figures S4.** Branch lengths across datasets and species tree inference methods (MIN4 datasets). A) Difference between discordance-based branch lengths estimated with ASTRAL-Pro (APro) and all other methods, normalized by APro branch length. B) Difference between site-based branch lengths for internal branches from the SCC dataset and all other datasets, normalized by SCC branch length. Site-based branch lengths are estimated using concatenated ML. C) Same as in panel C, but for terminal branches. Colors represent different filtering methods, and each row is a different branch. SCC=single-copy clusters; LSD=lineage-specific duplicates; TSD=two-species duplicates; MO=monophyletic outgroup; MI=maximum inclusion; SE=subtree extraction; ONE=one paralogs.

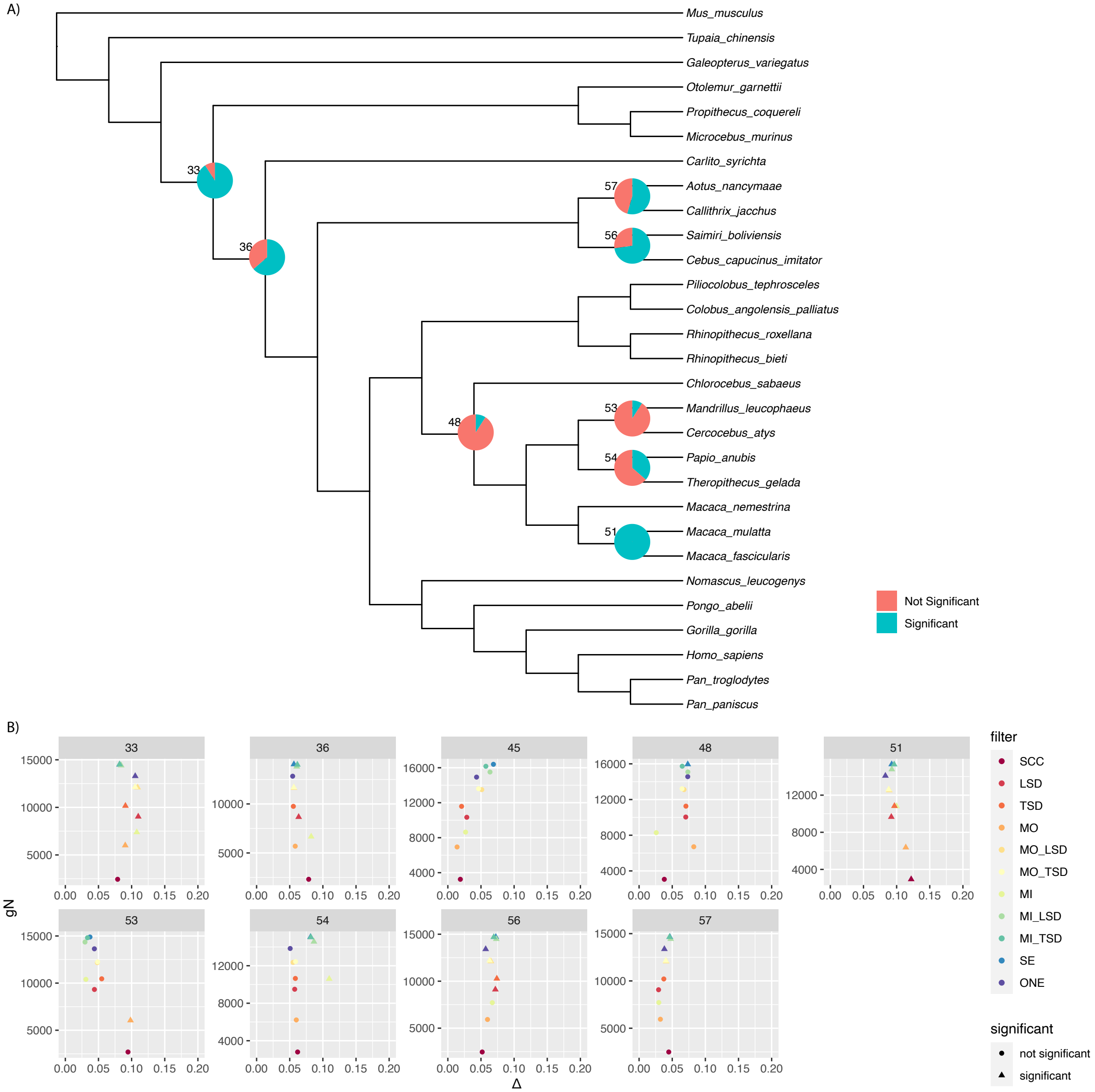


**Figure S5.** Results of introgression tests on MIN4 ML gene trees. A) Pie charts are shown for branches with any significant introgression tests. Numbers are node numbers. B) For all branches with some significant tests, we show the number of informative genes versus Δ. Observations are colored by filtering method, and shapes indicate whether a particular test was significant. SCC=single-copy clusters; LSD=lineage-specific duplicates; TSD=two-species duplicates; MO=monophyletic outgroup; MI=maximum inclusion; SE=subtree extraction; ONE=one paralogs.

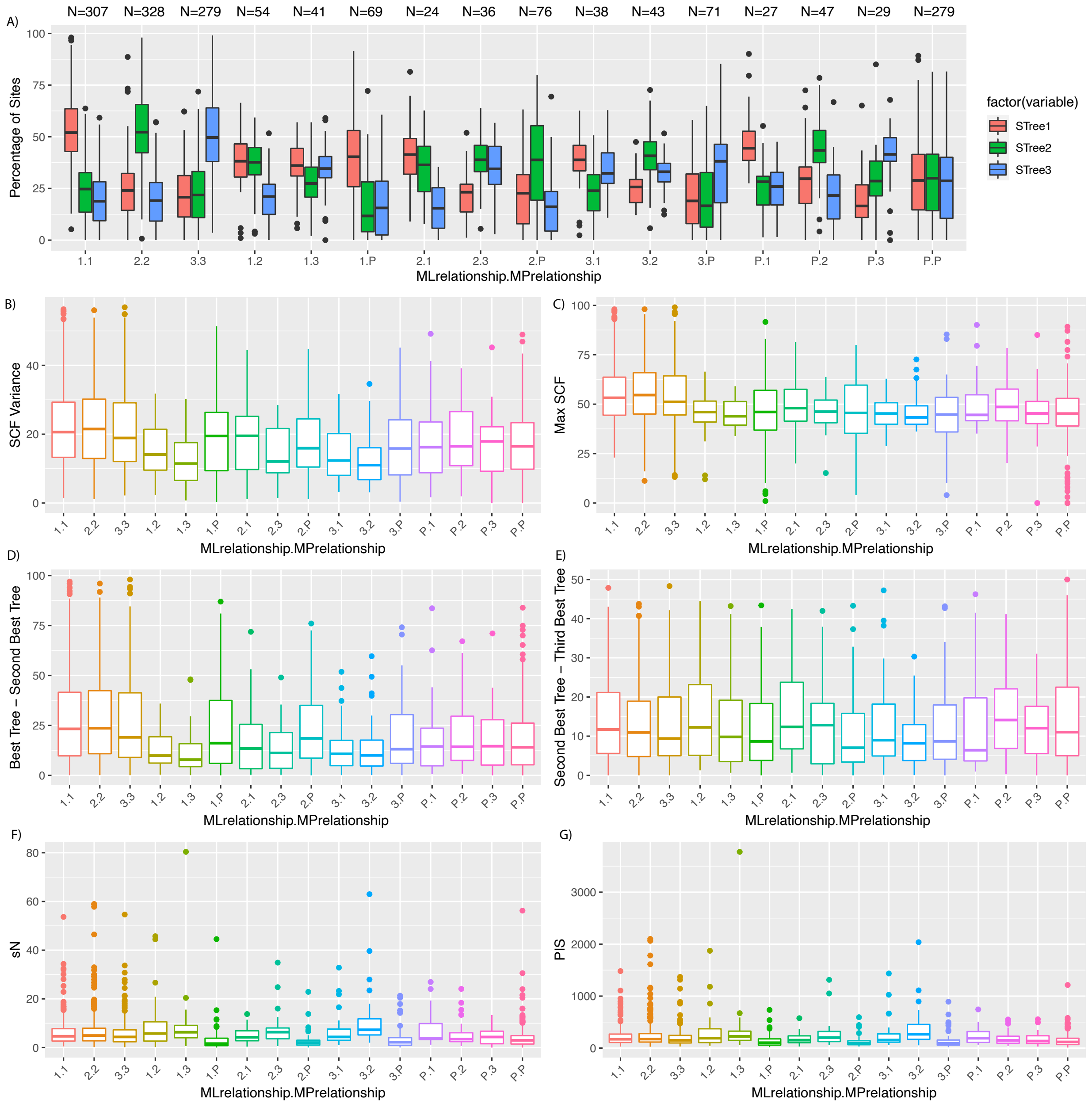


**Figure S6.** Comparisons among ML and MP gene trees with respect to relationships among the Platyrrhini (branch 57; Figure 3a). Trees 1-3 correspond to the trees shown in Figure 4 of the main text. Each category on the x-axis is a set of gene trees, divided by the tree supported by ML and MP. The tree preferred under ML inference is listed first, followed by a period, and then the tree preferred under MP inference. The tree labeled "P" represents cases where one of the three clades was not monophyletic, so the tree topology did not match one of the ones shown in Figure 4. The number of gene trees in each category is given in panel A. A) Percentage of decisive sites supporting each tree. B) Variance of sCF scores within genes. C) Maximum sCF within a gene. D) Maximum sCF minus second highest sCF within a gene. E) Second highest sCF minus lowest sCF within a gene. F) Number of sites informative about this branch ("decisive") for IQ-Tree sCF calculations. G) Number of Parsimony Informative Sites (PIS) in the alignment. Results from MIN27 datasets.

Appendix A: ***Alternative MI datasets***

To explore the effects of the branch-length cutoff used in the MI approach, we additionally considered a long (10 substitutions per site; 5000 changes for MP gene trees) threshold. Additionally, to explore the effects of including multiple gene trees per cluster (after tree decomposition), we generated datasets from the MI datasets where we randomly sampled a single gene tree per cluster. The results using the longer threshold and randomly sampling a single gene per cluster did not differ qualitatively from those presented in the main text, and are thus reported here. In general, using a longer branch length cutoff lead to very small decreases in the number of genes available (Table A1). Of course, sampling a single gene tree per cluster led to fewer genes and fewer informative sites (Table A1; Figure A1). Species trees were identical to those estimated from the original MI datasets. Gene and site concordance factors (Figures A2–A5), branch length estimates (Figures A6-A7), and introgression test results (Figures A8-A9) were similar to those from the original MI datasets.

| Filter | MIN4 | | MIN27 | |
|---|---|---|---|---|
| | Gene families | Gene copies | Gene families | Gene copies |
| *Maximum Inclusion* | 27880 | 331990 | 4849 | 137733 |
| *Maximum Inclusion (LSD)* | 22360 | 464224 | 11479 | 327434 |
| *Maximum Inclusion (TSD)* | 21793 | 473000 | 12046 | 343652 |
| *Maximum Inclusion (LONG)* | 27900 | 332233 | 4856 | 137962 |
| *Maximum Inclusion (LONG; LSD)* | 22362 | 464306 | 11483 | 327567 |
| *Maximum Inclusion (LONG; TSD)* | 21795 | 473076 | 12049 | 343758 |
| *ONE-Maximum Inclusion* | 17303 | 254286 | 4722 | 134168 |
| *ONE-Maximum Inclusion (LSD)* | 18467 | 403399 | 10486 | 299079 |
| *ONE-Maximum Inclusion (TSD)* | 18477 | 413779 | 10978 | 313146 |
| *ONE-Maximum Inclusion (LONG)* | 17310 | 254477 | 4729 | 134386 |
| *ONE-Maximum Inclusion (LONG; LSD)* | 18468 | 403196 | 10489 | 299155 |
| *ONE-Maximum Inclusion (LONG; TSD)* | 18478 | 414020 | 10980 | 313200 |

**Table A1.** Number of gene trees and gene copies included with different filtering approaches. LONG indicates datasets for which a long threshold was used for the MI filtering. LSD and TSD indicate when lineage-specific and both lineage-specific and two-species duplicates were trimmed. The ONE- prefix for MI datasets indicates datasets for which a single randomly selected gene tree per gene family was retained for downstream inference. The MIN4 dataset required a minimum of 4 taxa (out of 29 total), while the MIN27 dataset required a minimum of 27 taxa.

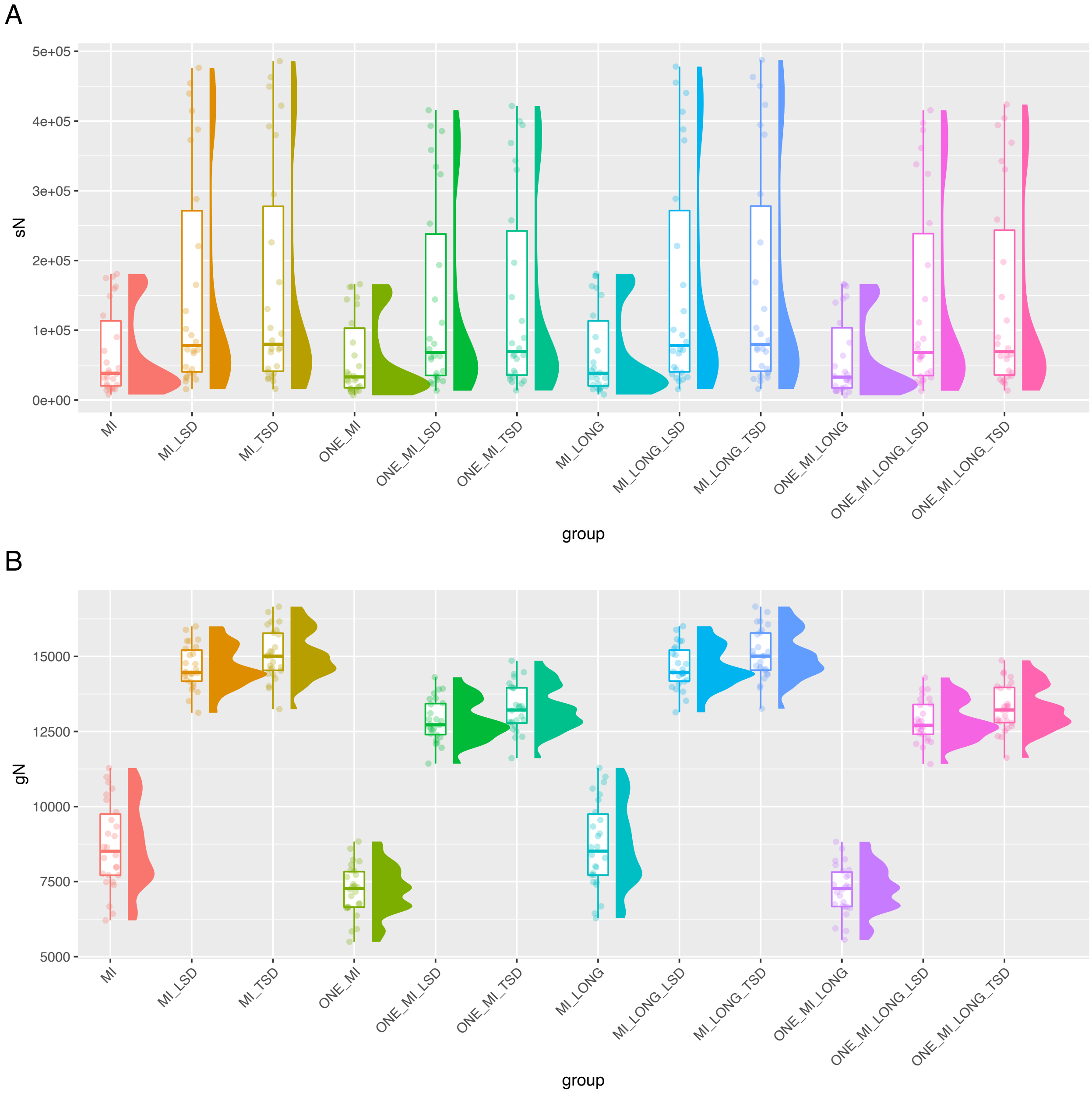


**Figure A1.** Numbers of informative genes and sites across datasets using the MIN27 MI datasets. A) The distribution of the number of decisive sites (across branches) as calculated in IQ-Tree. Decisive sites are defined in Minh et al. (2020). B) The distribution of the number of decisive gene trees (across branches) as calculated in IQ-Tree. Decisive gene trees are defined in Minh et al. (2020). ONE_MI = Maximum Inclusion with a single gene tree per cluster; MI_LONG=Maximum Inclusion with branch length cutoff of 10 substitutions per site; LSD=Lineage-Specific Duplicates trimmed; TSD=Two-Species Duplicates trimmed.

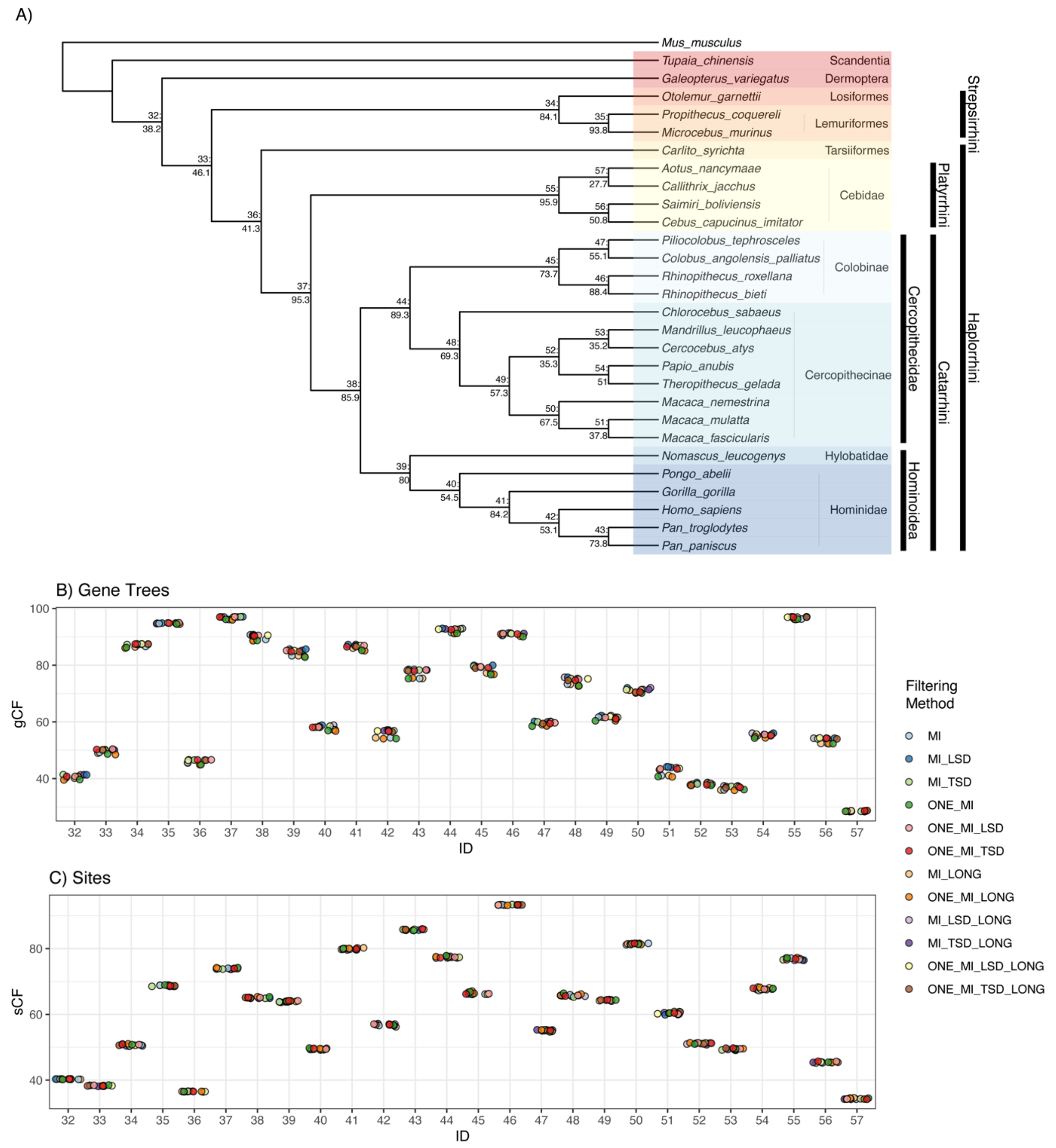


**Figure A2**. Gene (gCF) and site (sCF) concordance factors among primate datasets using ML gene trees (MIN4) and additional MI filtering. A) Primate phylogeny from ASTRAL-III using the ML gene trees (all input datasets give the same topology). Nodes show Node ID: gCF values from the SCC dataset. B) Distribution of gCF values across datasets. C) Distribution of sCF values across datasets. Node IDs correspond to the numbers displayed on the tree in panel A. LSD=lineage-specific duplicates; TSD=two-species duplicates; MI=maximum inclusion; Prefix 'ONE'=MI datasets with one gene tree sampled per cluster; Suffix 'LONG' MI datasets with longer branch length cutoff (10 substitutions per site).

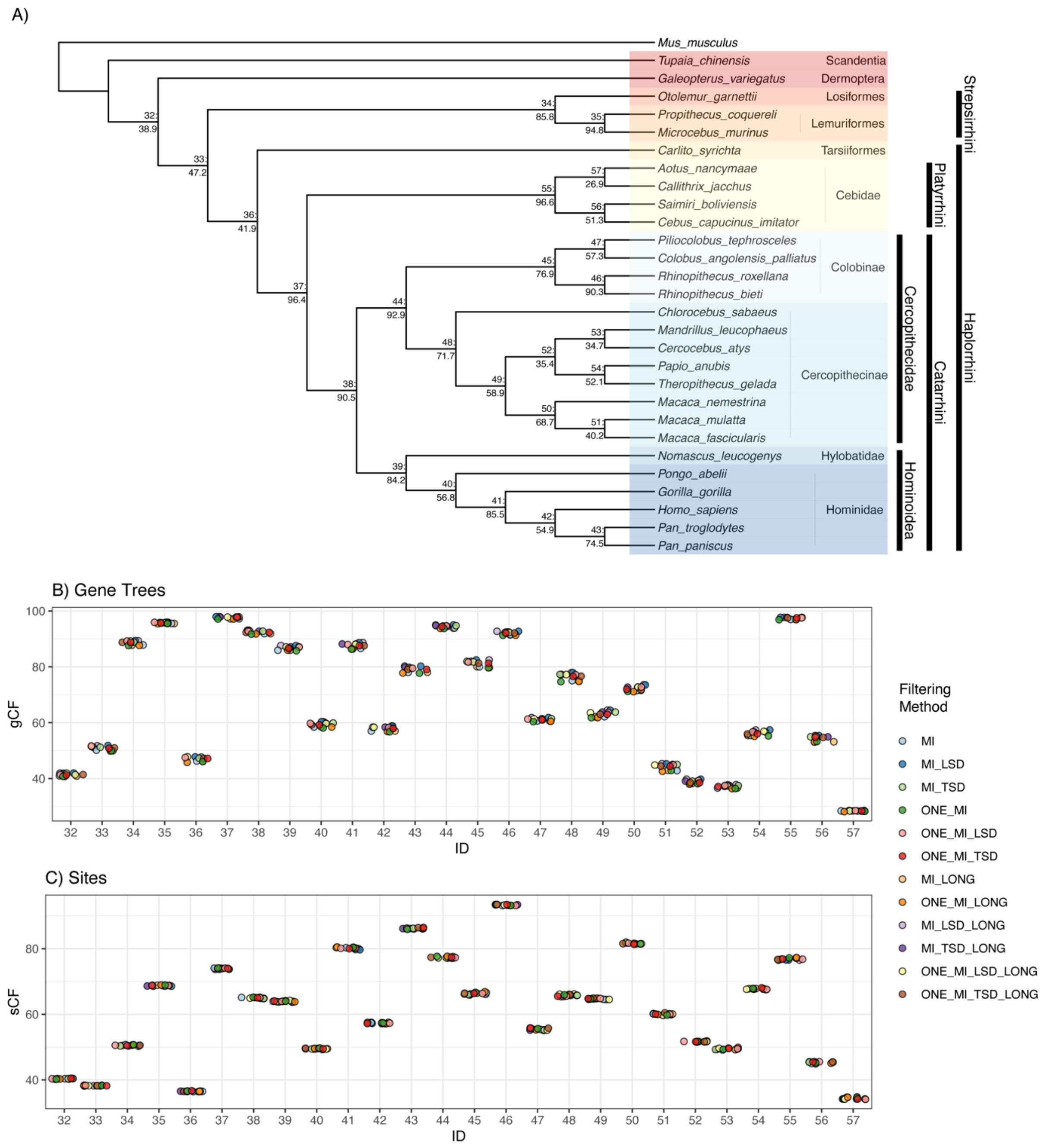


**Figure A3**. Gene (gCF) and site (sCF) concordance factors among primate datasets using ML gene trees (MIN27) and additional MI filtering. A) Primate phylogeny from ASTRAL-III using the ML gene trees (all input datasets give the same topology). Nodes show Node ID: gCF values from the SCC dataset. B) Distribution of gCF values across datasets. C) Distribution of sCF values across datasets. Node IDs correspond to the numbers displayed on the tree in panel A. LSD=lineage-specific duplicates; TSD=two-species duplicates; MI=maximum inclusion;. Prefix ‘ONE’=MI datasets with one gene tree sampled per cluster; Suffix ‘LONG’ MI datasets with longer branch length cutoff (10 substitutions per site).

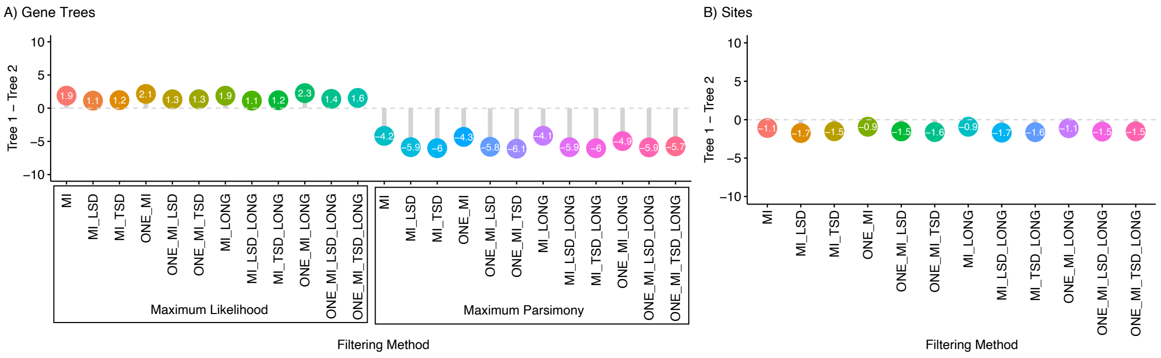


**Figure A4.** Alternative resolutions of Platyrrhini relationships using MIN4 datasets with additional MI filtering. A) The percentage of gene trees supporting Tree 1 minus the percentage of gene trees supporting Tree 2 for ML and MP gene trees across datasets. B) The percentage of sites supporting Tree 1 minus the percentage of sites supporting Tree 2 across datasets. LSD=lineage-specific duplicates; TSD=two-species duplicates; MI=maximum inclusion; Prefix ‘ONE’=MI datasets with one gene tree sampled per cluster; Suffix ‘LONG’ MI datasets with longer branch length cutoff (10 substitutions per site).

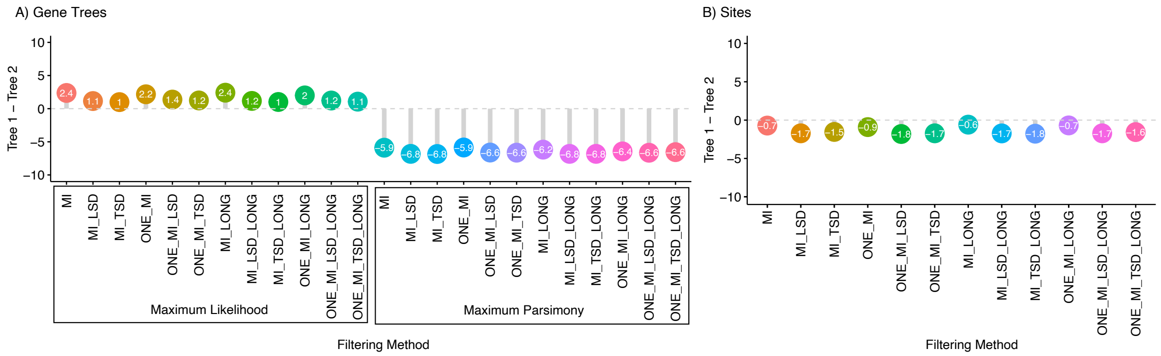


**Figure A5.** Alternative resolutions of Platyrrhini relationships using MIN27 datasets with additional MI filtering. A) The percentage of gene trees supporting Tree 1 minus the percentage of gene trees supporting Tree 2 for ML and MP gene trees across datasets. B) The percentage of sites supporting Tree 1 minus the percentage of sites supporting Tree 2 across datasets. LSD=lineage-specific duplicates; TSD=two-species duplicates; MI=maximum inclusion; Prefix 'ONE'=MI datasets with one gene tree sampled per cluster; Suffix 'LONG' MI datasets with longer branch length cutoff (10 substitutions per site).

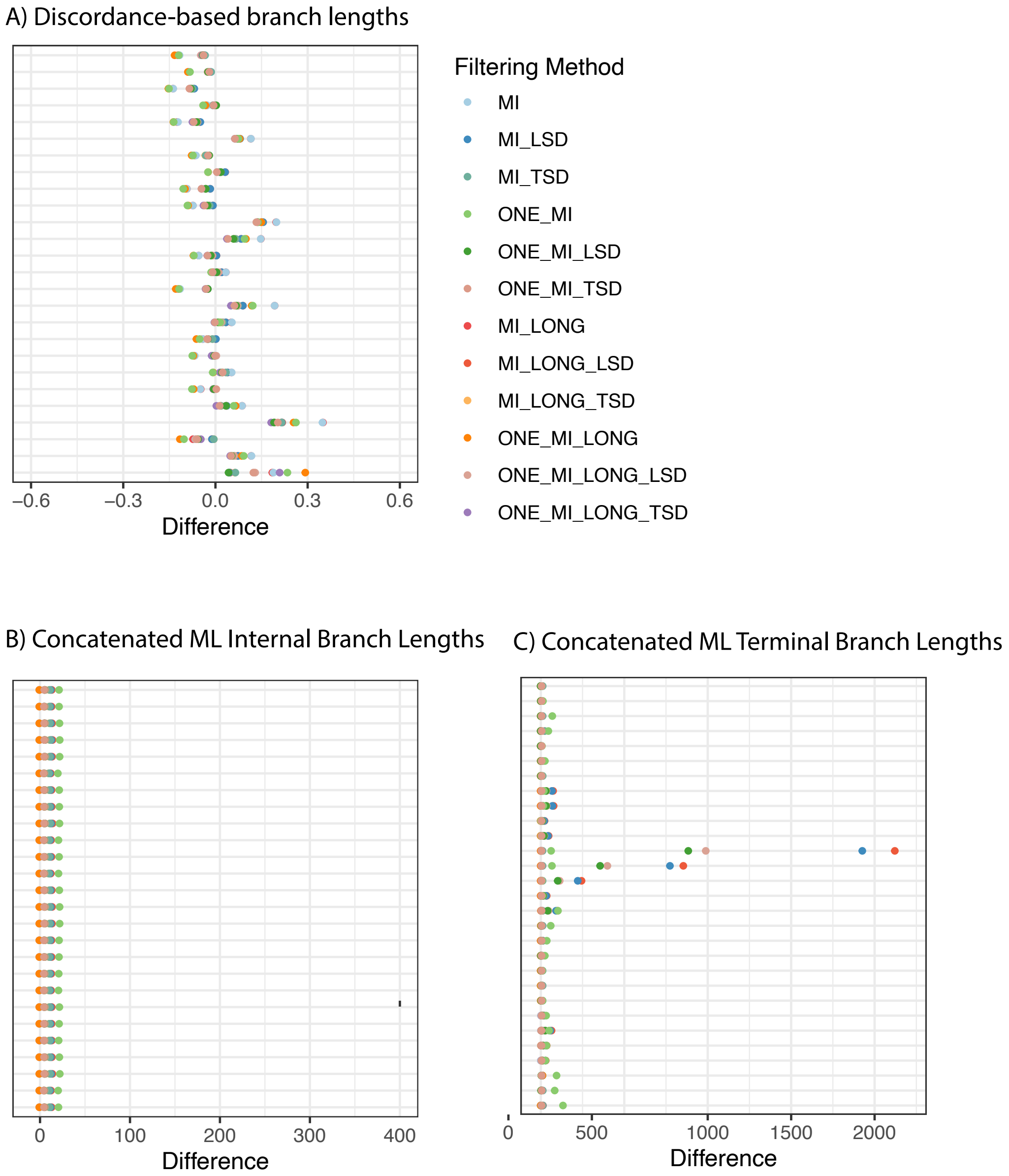


**Figures A6.** Branch lengths across datasets and species tree inference methods using MIN4 datasets and additional MI filtering. A) Difference between discordance-based branch lengths estimated with ASTRAL-Pro (APro) and all other methods, normalized by APro branch length. B) Difference between site-based branch lengths for internal branches from the SCC dataset and all other datasets, normalized by SCC branch length.  C) Same as in panel C, but for terminal branches. Colors represent different filtering methods, and each row is a different branch. LSD=Lineage-Specific Duplicates; TSD=Two-Species Duplicates; MI= Maximum Inclusion branch-cutting; Prefix 'ONE'=MI datasets with one gene tree sampled per cluster; Suffix 'LONG' MI datasets with longer branch length cutoff (10 substitutions per site)..

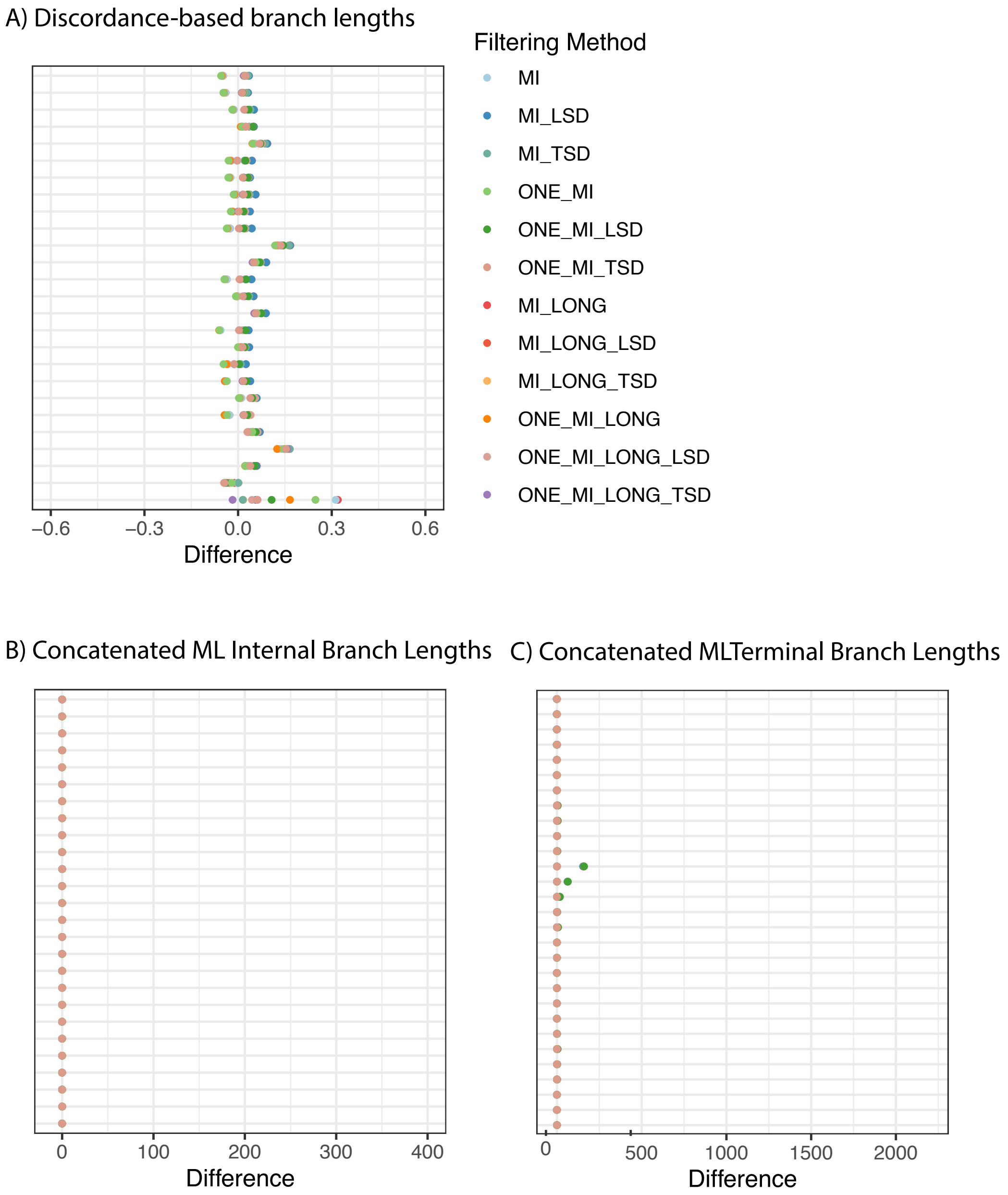


**Figures A7.** Branch lengths across datasets and species tree inference methods using MIN27 datasets and additional MI filtering. A) Difference between discordance-based branch lengths estimated with ASTRAL-Pro (APro) and all other methods, normalized by APro branch length. B) Difference between site-based branch lengths for internal branches from the SCC dataset and all other datasets, normalized by SCC branch length. C) Same as in panel C, but for terminal branches. Colors represent different filtering methods, and each row is a different branch. LSD=Lineage-Specific Duplicates; TSD=Two-Species Duplicates; MI= Maximum Inclusion branch-cutting; Prefix 'ONE'=MI datasets with one gene tree sampled per cluster; Suffix 'LONG' MI datasets with longer branch length cutoff (10 substitutions per site).

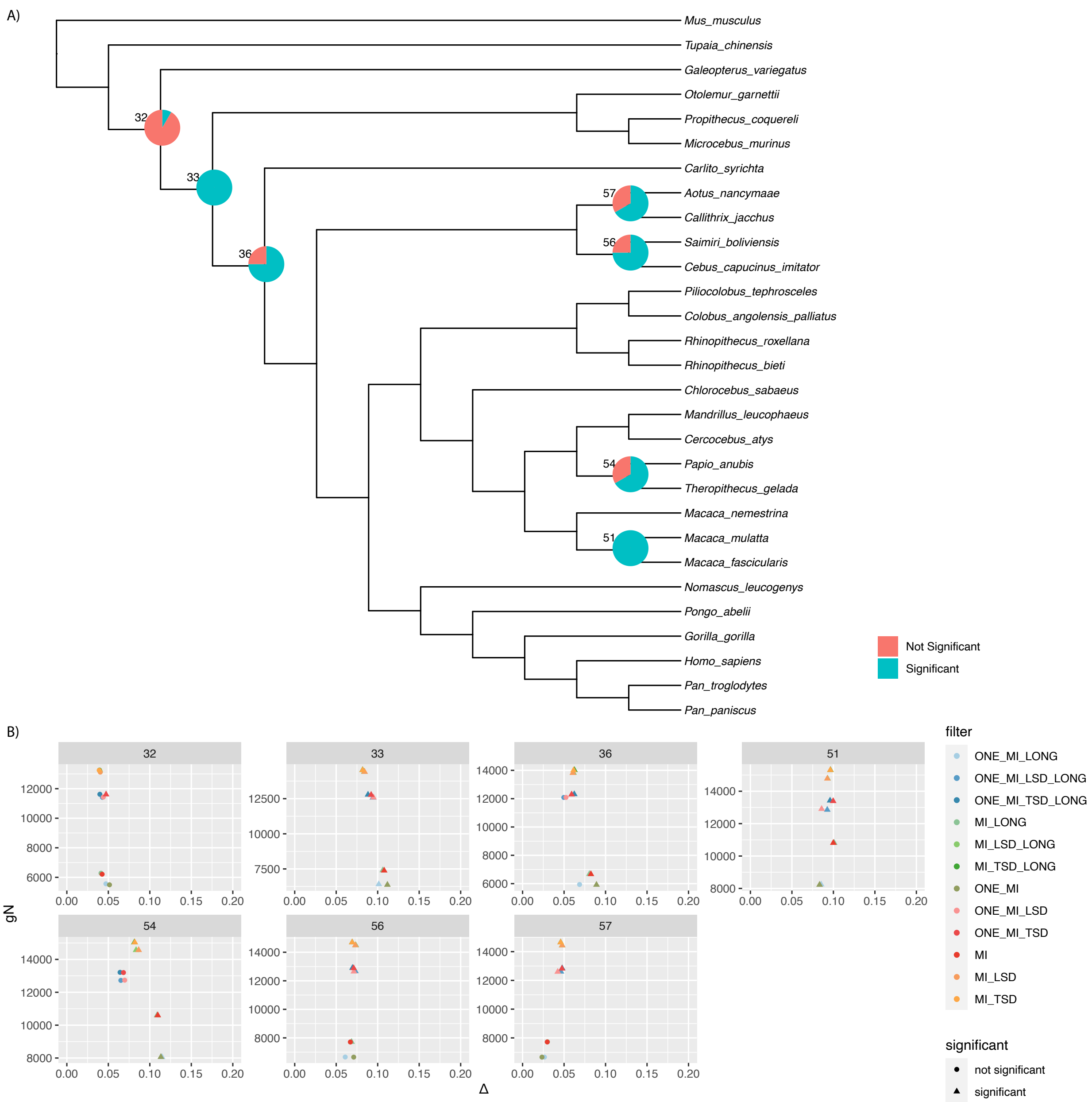


**Figure A8.** Results of introgression tests on MI gene trees using MIN4 datasets. A) Pie charts are shown for branches with any significant introgression tests. Numbers are node numbers. B) For all branches with some significant tests, we show the number of informative genes versus Δ. Observations are colored by filtering method, and shapes indicate whether a particular test was significant. LSD=Lineage-Specific Duplicates; TSD=Two-Species Duplicates; MI= Maximum Inclusion; Prefix 'ONE'=MI datasets with one gene tree sampled per cluster; Suffix 'LONG' MI datasets with longer branch length cutoff (10 substitutions per site).

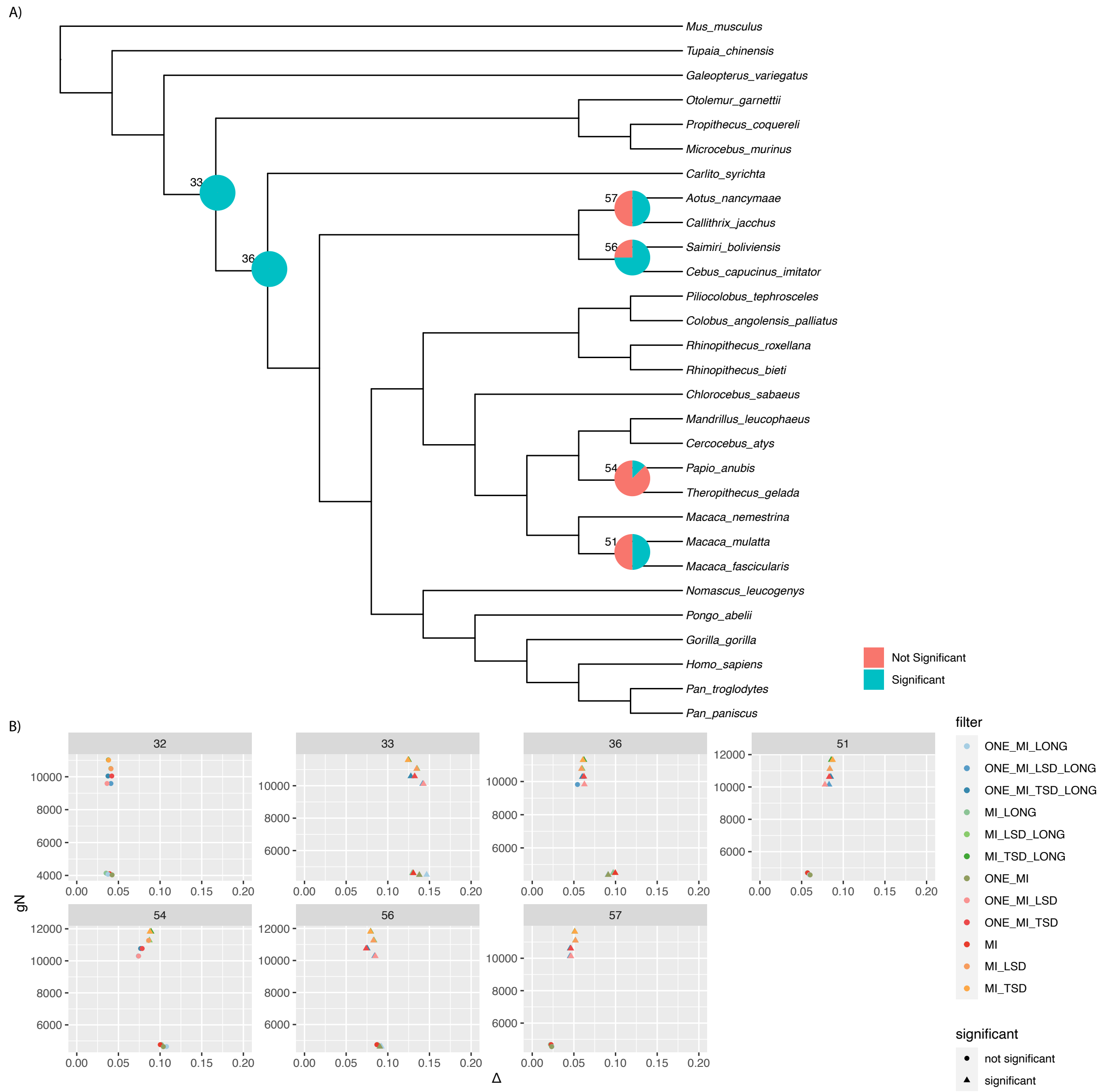


**Figure A9.** Results of introgression tests on MI gene trees using MIN27 datasets. A) Pie charts are shown for branches with any significant introgression tests. Numbers are node numbers. B) For all branches with some significant tests, we show the number of informative genes versus Δ. Observations are colored by filtering method, and shapes indicate whether a particular test was significant. LSD=Lineage-Specific Duplicates; TSD=Two-Species Duplicates; MI= Maximum Inclusion; Prefix ‘ONE’=MI datasets with one gene tree sampled per cluster; Suffix ‘LONG’ MI datasets with longer branch length cutoff (10 substitutions per site).

Appendix B: ***Analyses of non-primate datasets***

To explore the impacts of varying the number of species and genes and the depth of divergence, we analyzed five additional datasets from across the eukaryotic tree of life, focusing on inferences of species tree topologies.

***Fungi-16***

We reanalyzed gene families from 16 fungi species from Rasmussen and Kellis (2012) available from http://compbio.mit.edu/dlcoal/. These data consist of 5304 gene families present in a minimum of four taxa, including 3565 single-copy clusters (Table B1), and the deepest divergences are estimated to have occurred ~ 180 myr. We found that inferences of species tree topologies were largely consistent when gene-tree based methods (i.e., ASTRAL-III and ASTRID) were used, and that these topologies differed from those inferred when concatenated ML and concatenated MP were used (Figure B1). However, there was a single exception. The tree inferred from All Paralogs using ASTRAL-III differed from other ASTRAL-III and ASTRID trees at two nodes that have been considered contentious in previous studies, but matched the tree from Rasmussen and Kellis (2012) (Figure B1). *Ashbya gossypii* was inferred as sister to *Kluyveromyces lactis*, rather than *K. waltii*, and the placement of *Saccharomyces castellii* differed. The concatenated ML and MP trees differed from the ASTRAL-III and ASTRID trees at a single node by placing *Candida guillermondii* sister to *C. lusitaniae* rather than as sister to *Debaryomyces hansenii*. With the exception of the All Paralogs tree, differences were more pronounced across species tree inference methods than across datasets (Figure 7; Figure B2).

***Fungi-60***

We reanalyzed gene families from 60 fungi species extracted from the PhylomeDB database (Huerta-Cepas et al., 2014; Morel et al., 2022). These data consist of 5594 gene families present in a minimum of four taxa, including 1361 single-copy clusters (Table B2). We found that tree inference was largely consistent within methods (i.e., ASTRAL-III trees tend to be highly similar to other ASTRAL-III trees regardless of which dataset was used) with a single notable exception (Figure 7; Figure B3). The tree reconstructed from the All Paralogs dataset using ASTRAL-III differed substantially from other trees (Figure B3). In an attempt to better understand this, we also inferred a species tree from All Paralogs in FASTRAL. We found that the tree reconstructed by FASTRAL was much more similar to other inferred trees (Figure B3). We calculated the quartet scores for the ASTRAL-III tree inferred from All Paralogs, the ASTRAL-Pro tree, and the FASTRAL tree inferred from All Paralogs tree using ASTRAL-III and the All Paralogs dataset and found that the quartet score was higher for the ASTRAL-Pro tree (12,425,072,354) and the FASTRAL tree (12,425,305,589) than for the ASTRAL-III tree (12,380,755,710). This suggests that this aberrant result arose due to some aspect of the search space construction in ASTRAL-III, rather than due to anything inherent to the dataset. The tree inferred most often using ASTRAL-III, ASTRID, and ASTRAL-Pro was identical to the tree inferred in Morel et al. (2022) using ASTRAL-Pro. Maximum Parsimony trees differed from those inferred using all other methods. Overall, differences were again more pronounced across species tree inference methods than across datasets (Figure 7; Figure B3), with the exception of the apparently aberrant ASTRAL-III tree inferred from All Paralogs.

### *Vertebrates-22*

We reanalyzed gene families from 22 vertebrate species extracted from the PhylomeDB database (Huerta-Cepas et al., 2014; Morel et al., 2022). These data consist of 17734 gene families present in a minimum of four taxa, including 2989 single-copy clusters (Table B3). Given our computational constraints (94 hours, 500 Gb) neither ASTRAL-III nor FASTRAL could complete on the All Paralogs dataset. We found that all trees except the tree inferred from the One Paralogs dataset using concatenated ML were highly similar (Figure 7; Figure B4; Figure B5). The most commonly inferred tree was shared across inferences in ASTRAL-III (SE, MI, One Paralogs), ASTRAL-Pro, and ASTRID (SCC, LSD, TSD, SE, MI) and matched the tree inferred in Morel et al. (2022) using Astral-Pro, MiniNJ (Morel et al. 2022), and SpeciesRax (Morel et al. 2022) (Figure B5).

### *Vertebrates-188*

We reanalyzed gene families from 188 vertebrate species extracted from the Ensembl Compara database (Zerbino et al. 2018; Morel et al. 2022). These data consist of 30003 gene families present in a minimum of four taxa, including 8933 single-copy clusters (Table B4). Given our computational constraints (94 hours, 500 Gb) neither ASTRAL-III nor FASTRAL could complete on the All Paralogs dataset. We ran FASTRAL instead of ASTRAL-III on the SE and One Paralogs datasets due to computational constraints, and we did not run concatenated MP or concatenated ML on any vertebrates-188 datasets due to computational constraints. We found that all trees were highly similar, irrespective of dataset or inference method (Figure 7; Figure B6).

### *Plants-23*

We reanalyzed gene families from 23 plant species extracted from the PhylomeDB database (Huerta-Cepas et al., 2014; Morel et al., 2022). These data consist of 19248 gene families present in a minimum of four taxa, including 566 single-copy clusters (Table B5). We ran FASTRAL instead of ASTRAL-III on the All Paralogs dataset due to computational constraints. FASTRAL, ASTRAL-III, ASTRID, and ML trees were highly similar, except for the ML tree inferred from the One Paralogs dataset (Figure 7; Figure B7; Figure B8). MP trees differed substantially from other trees (Figure 7; Figure B7). Amongst ASTRAL-III, ASTRID, and most ML trees the only differences were in the placement of *Theobroma* and *Fragaria* (Figure B7).

| Filter | Gene families | Gene copies |
|---|---|---|
| *Single-copy clusters (SCC)* | 3565 | 45733 |
| *Lineage-specific duplicates (LSD)* | 3820 | 48840 |
| *Two-species duplicates (TSD)* | 3892 | 49734 |
| *Maximum Inclusion (TSD)* | 9698 | 78542 |
| *Subtree Extraction (SE)* | 6231 | 76079 |
| *All Paralogs* | 5304 | 82218 |
| *One Paralogs* | 5304 | 69544 |

**Table B1.** Number of gene trees and gene copies included with different filtering approaches from the fungi-16 dataset. We required a minimum of 4 taxa.

| **Filter** | **Gene families** | **Gene copies** |
|---|---:|---:|
| *Single-copy clusters (SCC)* | 1361 | 30272 |
| *Lineage-specific duplicates (LSD)* | 1816 | 52103 |
| *Two-species duplicates (TSD)* | 1928 | 56810 |
| *Maximum Inclusion (TSD)* | 38860 | 345257 |
| *Subtree Extraction (SE)* | 12809 | 311354 |
| *All Paralogs* | 5594 | 387955 |
| *One Paralogs* | 5594 | 230199 |

**Table B2.** Number of gene trees and gene copies included with different filtering approaches from the fungi-60 dataset. We required a minimum of 4 taxa.

| Filter | Gene families | Gene copies |
| --- | --- | --- |
| *Single-copy clusters (SCC)* | 2989 | 46021 |
| *Lineage-specific duplicates (LSD)* | 3678 | 57931 |
| *Two-species duplicates (TSD)* | 3730 | 58547 |
| *Maximum Inclusion (TSD)* | 179050 | 1101600 |
| *Subtree Extraction (SE)* | 75537 | 1011898 |
| *All Paralogs* | 17734 | 1456788 |
| *One Paralogs* | 17734 | 325246 |

**Table B3.** Number of gene trees and gene copies included with different filtering approaches from the vertebrates-22 dataset. We required a minimum of 4 taxa.

| **Filter** | **Gene families** | **Gene copies** |
|---|---:|---:|
| *Single-copy clusters (SCC)* | 8933 | 163102 |
| *Lineage-specific duplicates (LSD)* | 12006 | 353133 |
| *Two-species duplicates (TSD)* | 13285 | 459833 |
| *Maximum Inclusion (TSD)* | 247393 | 3237227 |
| *Subtree Extraction (SE)* | 55126 | 2956328 |
| *All Paralogs* | 30003 | 3713676 |
| *One Paralogs* | 30003 | 2547863 |

**Table B4.** Number of gene trees and gene copies included with different filtering approaches from the vertebrates-188 dataset. We required a minimum of 4 taxa.

| Filter | Gene families | Gene copies |
|---|---|---|
| *Single-copy clusters (SCC)* | 566 | 5281 |
| *Lineage-specific duplicates (LSD)* | 2085 | 28834 |
| *Two-species duplicates (TSD)* | 2987 | 43333 |
| *Maximum Inclusion (TSD)* | 248749 | 1030214 |
| *Subtree Extraction (SE)* | 75557 | 814577 |
| *All Paralogs* | 19248 | 1605532 |
| *One Paralogs* | 19248 | 341868 |

**Table B5.** Number of gene trees and gene copies included with different filtering approaches 1from the plants-23 dataset. We required a minimum of 4 taxa.

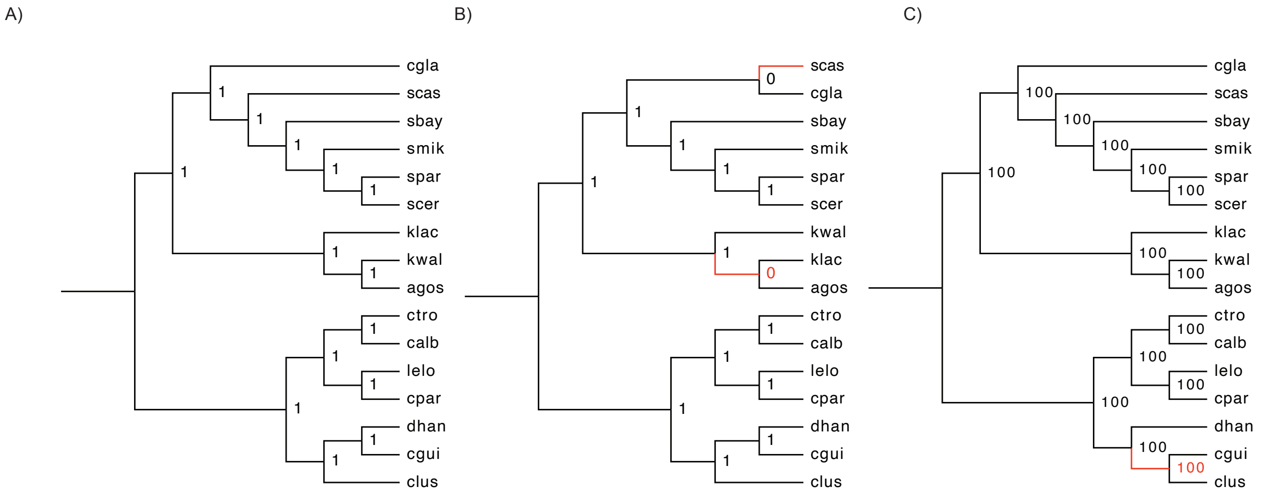


**Figure B1.** Results from species tree inference on the fungi-16 dataset. A) The tree inferred when running ASTRAL-III on all datasets except the All Paralogs dataset and ASTRID on all datasets. Node values are posterior probabilities inferred from the SCC dataset in ASTRAL-III. B) The tree inferred using ASTRAL-III on the All Paralogs dataset. Node values are posterior probabilities inferred from the All Paralogs dataset in ASTRAL-III. C) The tree inferred using concatenated ML and concatenated MP. Node values are bootstrap support values for the SCC dataset under concatenated ML inference. Branch lengths are not drawn to scale.

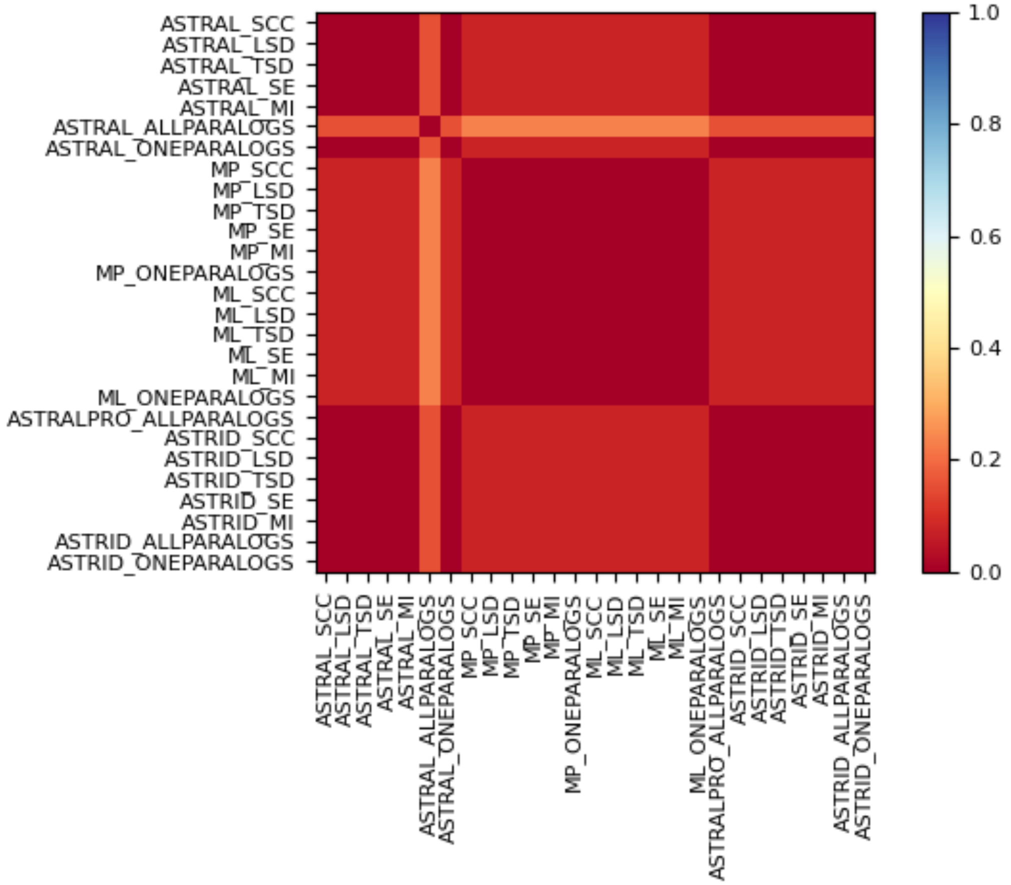


**Figure B2.** Normalized Robinson-Foulds distances between all species trees inferred from the fungi-16 dataset. SCC=single-copy clusters; LSD=lineage-specific duplicates; TSD=two-species duplicates; MI=maximum inclusion with two-lineage duplicates trimmed; SE=subtree extraction.

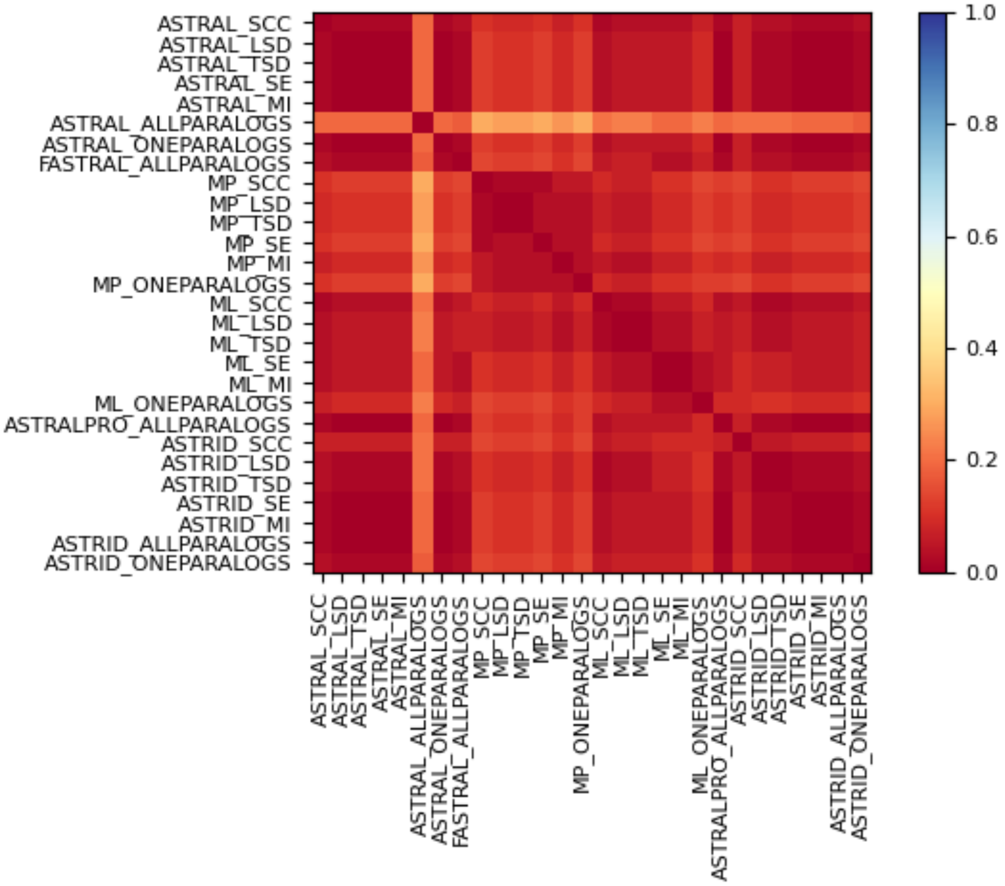


**Figure B3.** Normalized Robinson-Foulds distances between all species trees inferred from the fungi-60 dataset. SCC=single-copy clusters; LSD=lineage-specific duplicates; TSD=two-species duplicates; MI=maximum inclusion with two-lineage duplicates trimmed; SE=subtree extraction.

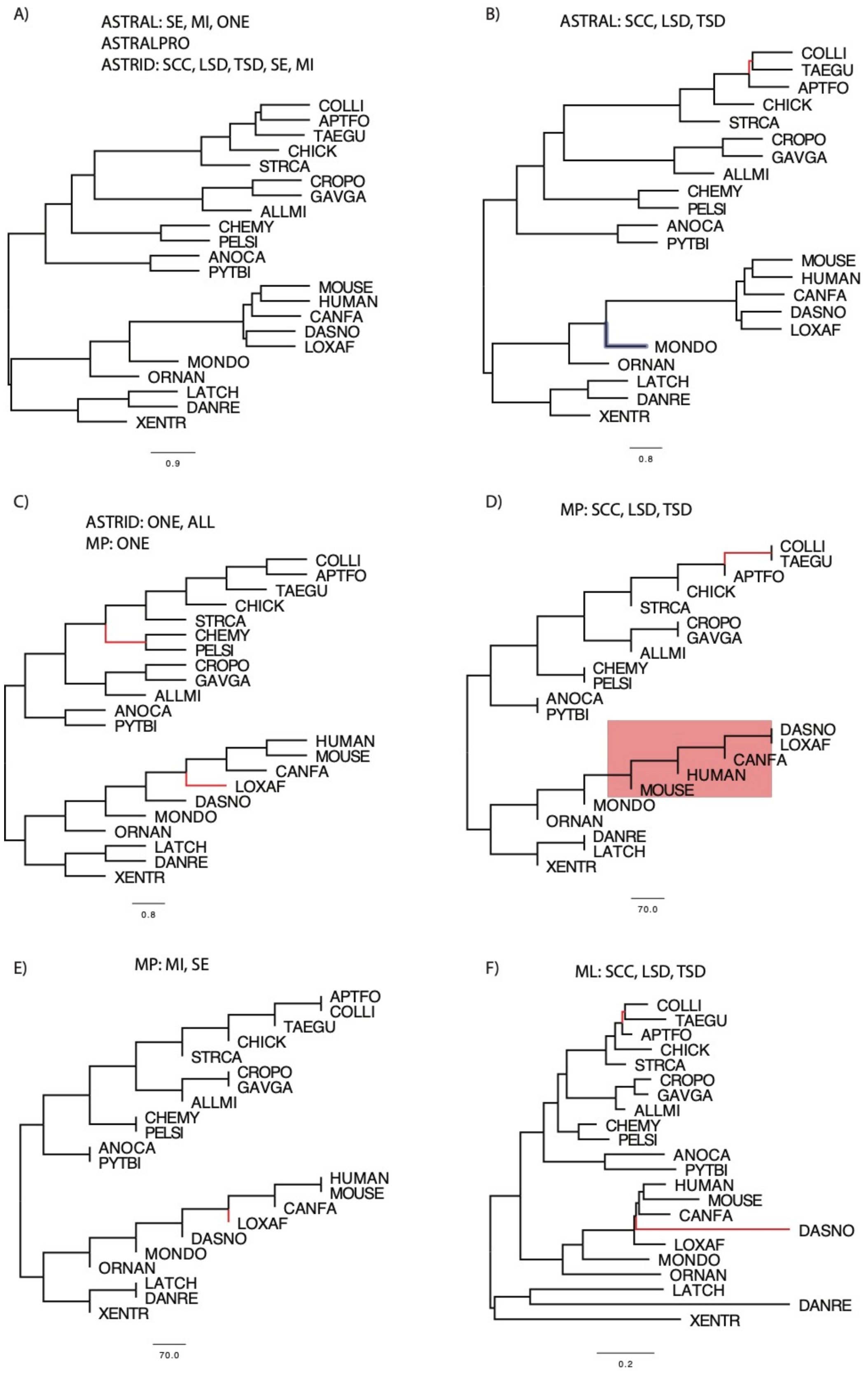


**Figure B3.** Trees inferred from the verebrates-21 dataset.

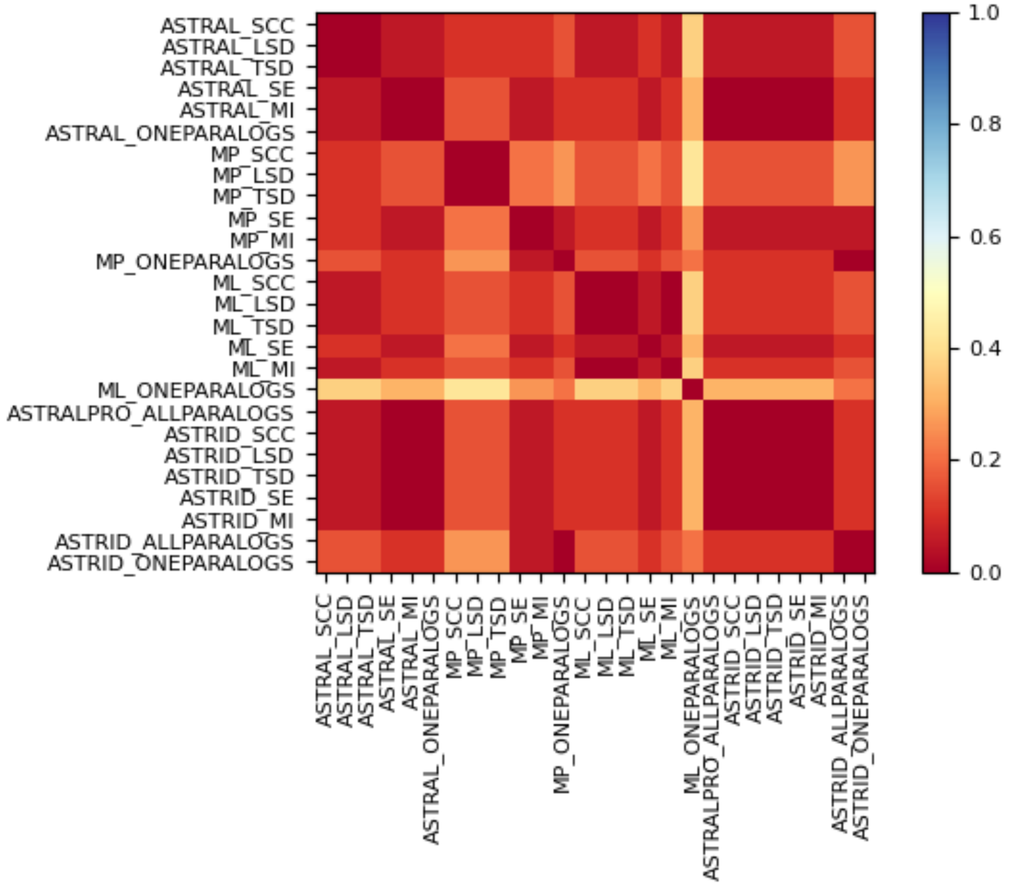


**Figure B4.** Normalized Robinson-Foulds distances between all inferred species trees inferred from the vertebrates-22 dataset. SCC=single-copy clusters; LSD=lineage-specific duplicates; TSD=two-species duplicates; MI=maximum inclusion with two-lineage duplicates trimmed; SE=subtree extraction.

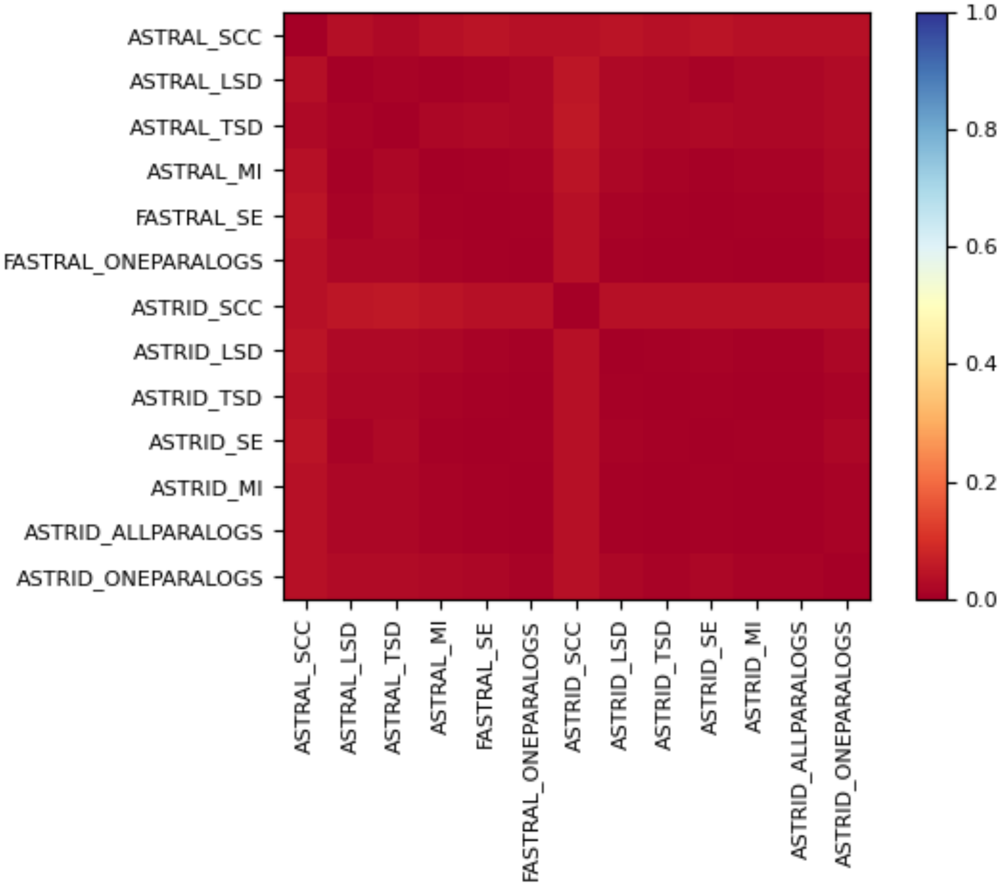


**Figure B5.** Normalized Robinson-Foulds distances between all inferred species trees inferred from the vertebrates-188 dataset. SCC=single-copy clusters; LSD=lineage-specific duplicates; TSD=two-species duplicates; MI=maximum inclusion with two-lineage duplicates trimmed; SE=subtree extraction.

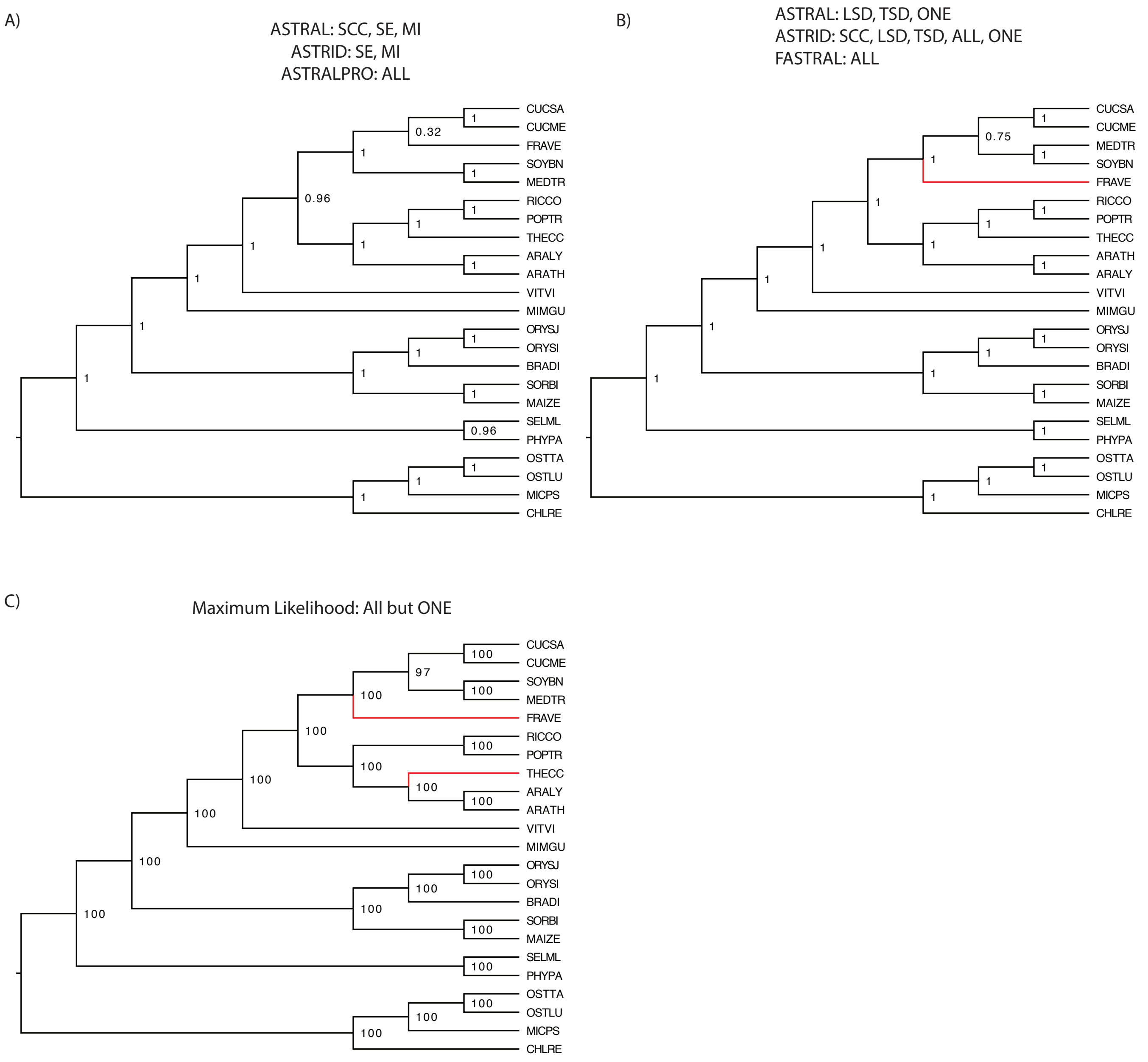


**Figure B6.** Results from species tree inference on the plants-23 dataset. A) The tree inferred when running ASTRAL-III on SCC, SE, and MI datasets and ASTRID on SE and MI datasets. Node values are posterior probabilities inferred from the single-copy orthologs dataset in ASTRAL-III. B) The tree inferred using ASTRAL-III on the LSD, TSD, and One Paralogs datasets, ASTRID on the SCC, LSD, TSD, All Paralogs, and One Paralogs datasets, FASTRAL on the All Paralogs dataset, and ASTRAL-Pro. Node values are posterior probabilities inferred from the TSD dataset in ASTRAL-III. C) The tree inferred using Maximum Likelihood on all datasets but the One Paralogs dataset. Node values are bootstrap support values for the SCC dataset. Branch lengths are not drawn to scale.

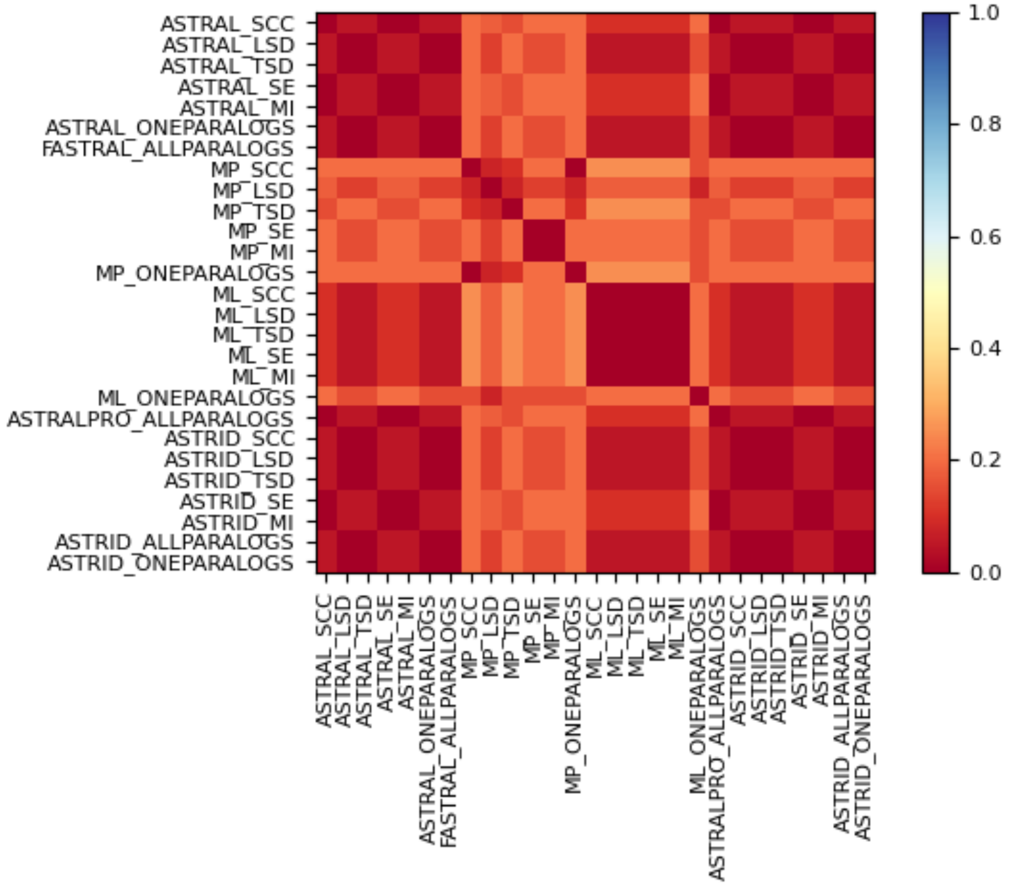


**Figure B7.** Normalized Robinson-Foulds distances between all inferred species trees inferred from the plants-23 dataset. SCC=single-copy clusters; LSD=lineage-specific duplicates; TSD=two-species duplicates; MI=maximum inclusion with two-lineage duplicates trimmed; SE=subtree extraction; ONE=one paralogs; ALL=all paralogs.

Zerbino DR, Achuthan P, Akanni W, Amode M, Barrell D, Bhai J, Billis K, Cummins C, Gall A,
